# Screening metatranscriptomes for ultrastable RNA secondary structures reveals hidden bacteriophages and novel capsid nanomaterials

**DOI:** 10.64898/2026.04.05.716407

**Authors:** Daniel A. Villarreal, Nino Makasarashvili, Aaryan Kapoor, Max Root, Matthew Campbell, Skylar Gibson, Conor Schiveley, Amineh Rastandeh, Sherry Baker, Sundharraman Subramanian, Uri Neri, Carolyn E. Mills, Katelyn McNair, Anca M. Segall, Uri Gophna, Kristin N. Parent, Rees F. Garmann

## Abstract

Metatranscriptomics has transformed our view of RNA bacteriophage diversity, revealing vast numbers of single-stranded RNA (ssRNA) phages whose protein capsids can be engineered for biotechnology applications. However, many ssRNA phages remain hidden from current detection methods, which require protein-level similarity to known phages. Here we show that RNA structure provides an additional signal for the detection of ssRNA phages in metatranscriptomes, including hidden phages missed by prior protein-based methods. By computationally folding each contig and screening for exceptionally stable RNA secondary structures, we find evidence of thousands of previously unrecognized phages encoding novel coat proteins. We express a library of 12,000 such coat proteins in E. coli and find that most assemble into nuclease-resistant capsids. We determine the 3D structure of one such capsid by cryo-electron microscopy and demonstrate that it can be disassembled and reassembled in vitro to package heterologous RNA—a key step toward repurposing these particles as RNA delivery vehicles. We compile the newly discovered ssRNA phages with previously known ones into a database that contains sequence and structural information for over 460,000 unique RNA molecules and over 100,000 distinct coat proteins, providing a comprehensive resource for microbiology and nanomaterials research.

## Introduction

Bacteriophages (“phages”) are the most abundant viruses on Earth, yet their diversity remains largely unexplored. Recent estimates suggest that over 10^8^ distinct phage lineages inhabit our planet (1), but less than 0.01% have been observed or studied. Of the phages that have been studied, we know far more about those with DNA genomes than those with RNA genomes (2). Expanding our knowledge of RNA-phages—particularly those with single-stranded (ss) RNA genomes—is important not only for basic research (3) but also for biotechnology, where their protein capsids are being engineered as nanoscale materials for molecular display and cargo delivery (4, 5).

Metatranscriptomics has greatly expanded our view of ssRNA phage diversity, revealing vast numbers of previously unknown phages. By searching environmental RNA sequencing data (“metatranscriptomes”) for contiguous RNA sequences (“contigs”) that encode phage-specific proteins—often an RNA-dependent RNA polymerase (“RdRp”)—researchers have identified new phages without ever culturing them or their bacterial hosts in the lab (6). For example, Neri et al. (7) identified thousands of new ssRNA phages by detecting their RdRps via sequence-similarity to those of known phages. Independent studies have applied similar approaches and identified additional phages (8, 9), supplying a wealth of potential capsid-based nanomaterials (10).

But many other ssRNA phages—namely, those lacking familiar RdRp sequences—remain hidden within existing metatranscriptomes. Detecting these hidden phages requires methods that go beyond sequence similarity. One approach is to incorporate structural information (11). Indeed, Hou et al. (12) recently used AI-guided protein-structure prediction to detect RdRps with the expected structure but little sequence similarity to known polymerases. While such protein-structure-based detection schemes have further expanded our view of phage diversity, they too have limitations: they cannot detect phages with entirely novel RdRps, or partial genomes with missing or incomplete RdRps but otherwise functional capsid genes.

To surmount these challenges, we exploit an orthogonal source of structural information—the secondary structure of the RNA. Like RdRps, phage RNAs adopt structures that are essential for replication (13). In principle, one could identify ssRNA phages in metatranscriptomes by predicting the RNA structure of each contig and then detecting phage-specific motifs. However, RNA structure prediction is notoriously inaccurate for sequences the length of phage genomes (14) and there have been no reports of broadly conserved structural motifs within the genomes of known phages (15). For these reasons, we instead focus on “genome-scale” (16) properties of RNA secondary structure, such as overall thermodynamic stability. Compared to motif detection, quantitation of thermodynamic stability is more robust to the inaccuracies of RNA structure prediction. And, as we demonstrate below, high thermodynamic stability is a broadly conserved property among known ssRNA phages, providing a signature that can be used to discover new ones.

Our approach is straightforward (see **Fig. 1, top row**): we apply RNA folding algorithms to previously published metatranscriptomes and keep only those contigs whose thermodynamic stability—defined using Z-scores, as detailed below—matches that of known phages. Within this pool of retained contigs, we find increased prevalence of previously identified phage genomes. In addition, we identify novel contigs encoding proteins with the canonical coat-protein fold, indicating the discovery of new phages. We consolidate these new phage genomes and their encoded coat proteins with those of previously known phages to build a database of sequence and structural information for over 460,000 distinct phage RNAs and 100,000 distinct coats.

**Fig. 1.**
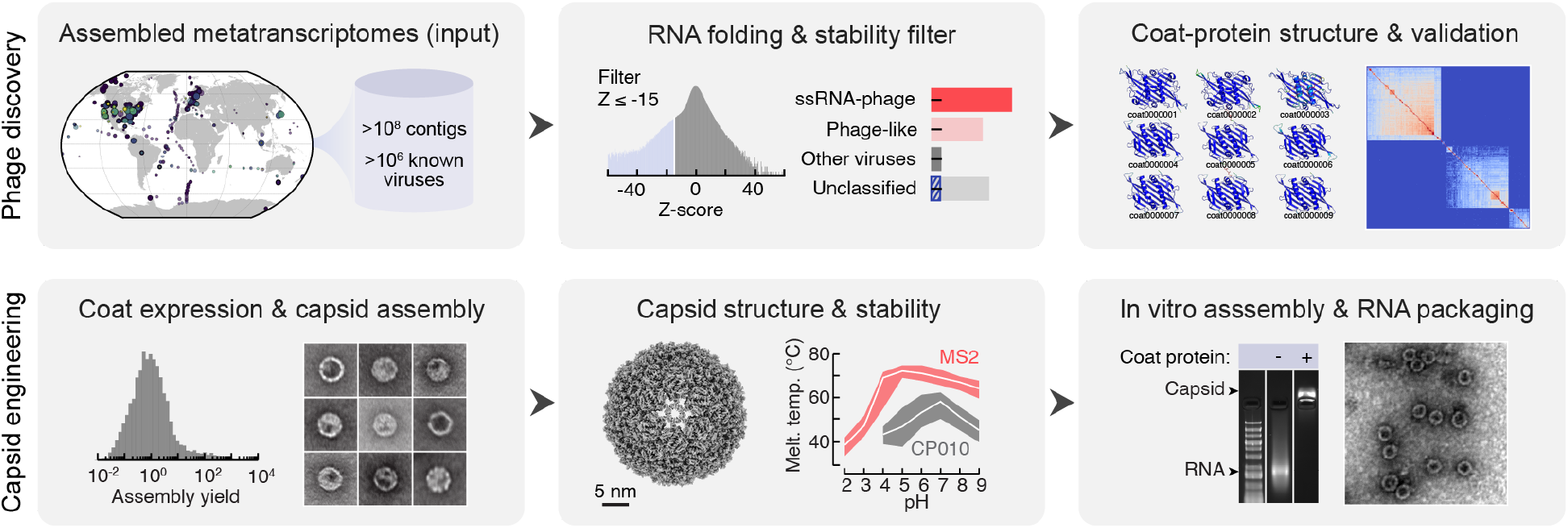
Our approach. **Top:** Starting from a previously reported metatranscriptome (left), we use RNA folding models to predict the secondary structure of each contig, retaining only those with exceptional thermodynamic stability (middle). The pool of retained contigs is enriched in known and novel phages, which encode a broad range of coat proteins (right). **Bottom:** We express a library of 12,000 distinct coat proteins in E. coli and measure their relative assembly yields (left). For one individual capsid, we determine its 3D structure and biophysical properties (middle), and show that it can package foreign RNA in in vitro assembly experiments (right).

To explore their potential as nanomaterials (**Fig. 1, bottom row**), we express 12,000 ssRNA phage coat proteins in E. coli and show that many readily assemble into nuclease-resistant capsids. For one such capsid, we determine its three-dimensional structure and demonstrate that it can be disassembled and reassembled around foreign RNA.

## Results and discussion

### The genomes of ssRNA phages adopt exceptionally stable secondary structures

In 1976, for the first time in history, researchers sequenced a complete genome: the 3,569-nucleotide single-stranded RNA genome of bacteriophage MS2 (17). Using early methods of RNA structure prediction, it was shown that MS2 RNA could adopt extensive secondary structure (17). Here, 50 years later, we have revisited the secondary structure of the MS2 genome using modern RNA folding algorithms (18).

Given an RNA sequence, RNA folding algorithms predict the most probable secondary structures by identifying sets of base pairs that minimize the free energy. Although the predicted base pairs often differ from experimentally determined ones, the minimum free energy (MFE) nonetheless provides a quantitative measure of structural stability that can be computed directly from sequence alone. Throughout this work, we have focused on the relative stability of predicted secondary structures rather than the structures themselves.

We used RNAfold (19) to compute the MFE of MS2 RNA and expressed the result as a Z-score relative to 1,000 random shuffles of the sequence (20) (see **Methods**). This analysis yielded a Z-score of -19, indicating that MS2 RNA has significantly higher thermodynamic stability than expected for a random sequence of equal length and nucleotide composition (**Fig. 2A, red**). To place this result into context, YehI, an E. coli messenger RNA of comparable length, has a Z-score of only -1 (**Fig. 2A, blue**), and 23S ribosomal RNA, a highly structured 2.9-kilobase transcript that forms part of the E. coli ribosome, has a Z-score of -5 (**Fig. 2A, gray**). Thus, MS2 RNA appears exceptionally stable, even relative to the most structured cellular RNAs.

**Fig. 2.**
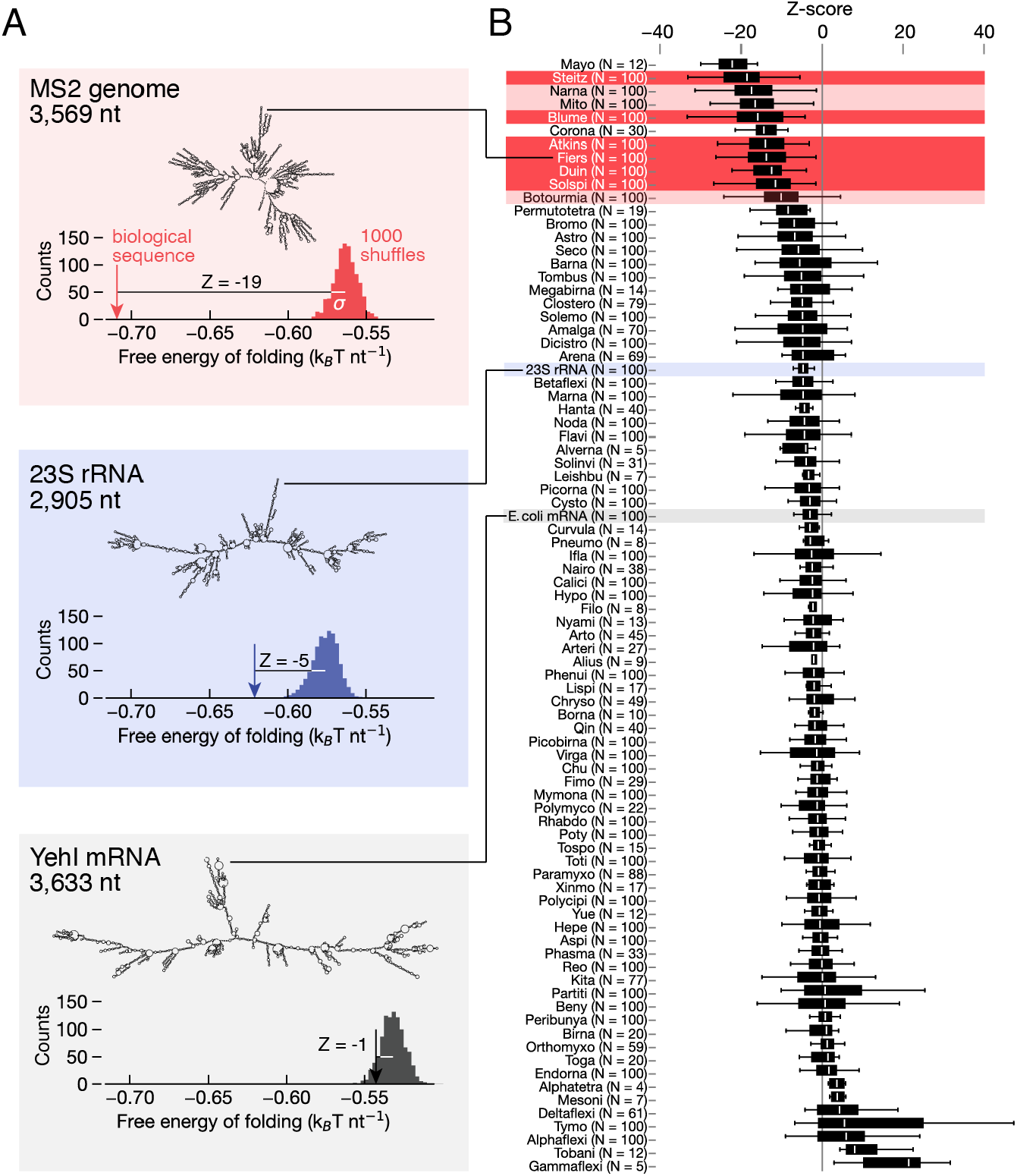
Quantifying thermodynamic stability of viral and non-viral RNA secondary structures. **A**. The MFE of MS2 RNA is 19 standard deviations lower than the mean MFE of 1000 random shuffles (Z=-19). Both the 23S rRNA and the mRNA of yehI are far less stably folded. **B**. Boxplots show Z-score distributions across RNA virus families, with boxes indicating medians and interquartile ranges, and bars indicating the 1st and 99th percentiles. ssRNA-phages (red) and phage-like viruses (light red) have among the most negative values.

To test whether this exceptional stability is unique to MS2, we extended our analysis to all families of RNA viruses listed in the Riboviria database (https://riboviria.org) (7). For each family, we randomly selected up to 100 complete genome sequences, computed their minimum free energies using RNAfold, and expressed the results as Z-scores (see **Methods**). When sorted by median Z-score, families of ssRNA phages consistently ranked among the most stable (**Fig. 2B, red boxes**). Also near the top of the list were families of phage-like viruses— including the narnaviruses, mitoviruses, and botourmiaviruses—which are thought to share recent ancestry with phages based on RdRp similarity (21) (**Fig. 2B, light red boxes**). By contrast, the median Z-scores of many families of eukaryotic ssRNA viruses fell closer to 0, similar to cellular mRNA (**Fig. 2B, gray box**).

These results show that ultrastable folding is a consistent and distinguishing property of ssRNA phages and their closest relatives. We suspect several selective pressures acted together to drive the evolution of such stable RNA structures, including pressures to form packaging signal stem-loops that promote capsid assembly (22), intramolecular duplexes that suppress deleterious plus-minus hybridization during replication (23), and structured regions that resist binding by CRISPR guide RNAs and other nucleic-acid surveillance pathways used in host-cell defense systems (24). Some of these pressures are unique to ssRNA phages, and might explain why their genomes are more stable than those of other viruses.

### Screening metatranscriptomes for stable structures enriches the prevalence of known phages and reveals hidden ones

We then asked if folding stability could be used to detect ssRNA phage genomes within metatranscriptomes. To test this idea, we computed the MFE Z-score for each contig in two recently published metatranscriptomic datasets, retained those contigs with Z ≤ -15, and then compared the prevalence of known phages before and after the Z-score filter. The first dataset that we analyzed, reported by Neri et al. (7), contained 4.7×10^6^ contigs that were classified by RdRp sequence, whereas the second dataset, reported by Hou et al. (12), contained 1.2×10^8^ contigs classified by RdRp structure. Because both of these datasets yielded qualitatively similar outcomes, we present our results for the Neri dataset below and provide the corresponding analysis of the Hou dataset as **Supporting Information**.

The distribution of Z-scores for the Neri dataset peaked at Z=0 with a tail extending toward negative values (**Fig. 3A**). Applying the Z ≤ -15 filter reduced the size of the dataset by two orders of magnitude to 4.0×10^4^ contigs but markedly enriched the prevalence of phage-related sequences. Before filtering, a vast majority of contigs were unclassified by prior RdRp-based analysis (98.0%), with only small fractions corresponding to ssRNA phages (0.4%), phage-like viruses (0.5%), or other viruses (1.1%) (**Fig. 3B, left**). After filtering, however, 24.5% of retained contigs were derived from ssRNA phages and 17.7% from phage-like viruses (**Fig. 3B, middle**), representing enrichments of ∼60-fold and ∼35-fold, respectively (**Fig. 3B, right**). As expected from our conservative Z-score threshold, nearly half of the annotated phage contigs in the original dataset (8,815 of 18,672, or 47%) had Z-scores above the cutoff and were not captured.

**Fig. 3.**
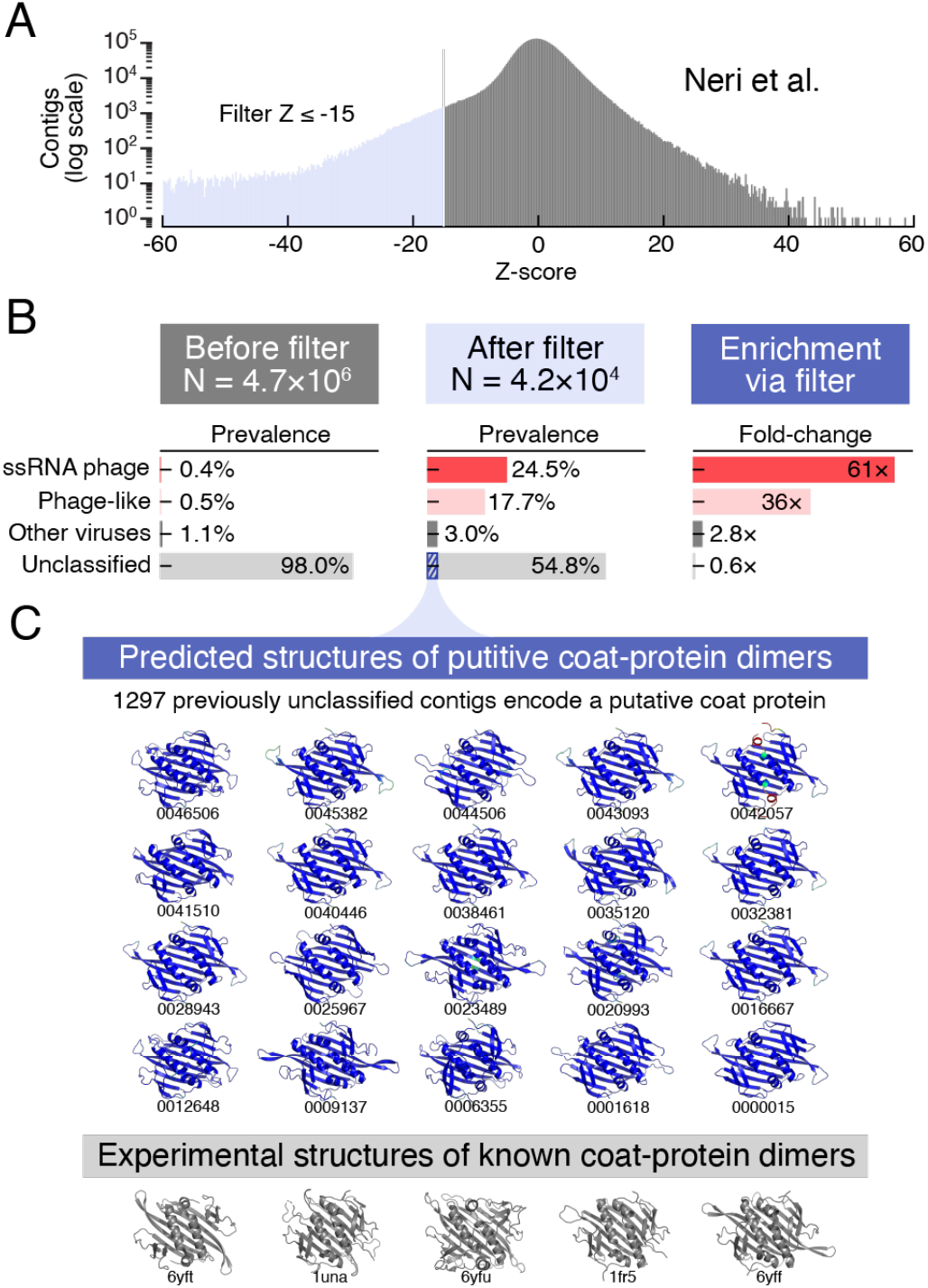
Filtering metatranscriptomes for ultrastable RNA secondary structures reveals both known and novel ssRNA-phages. **A**. The Z-score distribution of the Neri dataset (7). We filtered the dataset, retaining contigs with Z ≤ -15 (light blue). **B**. Prior to filtering, most of the contigs were unclassified (light gray), with relatively small fractions of ssRNA phages (red) and phage-like (light red) viruses. The prevalence of all other virus families was also small (dark gray). After filtering, however, the prevalence of ssRNA phage and phage-like viruses increased considerably. Furthermore, many of the previously unclassified contigs retained by the filter were found to encode proteins with the canonical ssRNA-phage coat-protein fold (blue stripes). **C. Top:** Predicted structures of several randomly selected coat-protein dimers. **Bottom:** Experimentally determined structures of known phage coat-protein dimers, for reference.

Despite this enrichment, most of the ultrastable contigs retained by our filter remained unclassified by prior RdRp-based analysis (54.8%), so we searched them for coat-protein genes as a hallmark of phage origin. This search identified 1,297 contigs encoding a protein predicted to adopt the canonical ssRNA-phage coat-protein fold (25) (see **Methods** and **Supporting Information, Fig. S1**). In the absence of a recognizable RdRp, the presence of a properly folded coat protein provides strong evidence that these ultrastable contigs derive from genuine phage genomes rather than unrelated structured RNAs. Together, these results show that our Z-score stability filter enriches not only known phage genomes but also uncovers previously unrecognized phage-derived contigs. Indeed, amino acid sequence analysis revealed that some of the new coats cluster with previously known examples, whereas others are entirely unique (**Supporting Information, Fig. S2**).

Next we turned to the Hou dataset. Applying the Z ≤ -15 filter reduced the size of the Hou dataset by over two orders of magnitude to 2.6×10^5^ contigs. Remarkably, this ultrastable population showed a ∼150-fold enrichment in the prevalence of known ssRNA phage and phage-like viruses and contained 10,219 previously unclassified phage-derived contigs encoding a characteristic coat-protein (see **Supporting Information, Fig. S3**). Many of these coat-encoding contigs contain fewer than 1,000 nucleotides—too few to encode a complete RdRp—which likely explains why they were missed by previous RdRp-detection methods (**Supplementary Information, Fig. S4**). Some of these contigs likely reflect incomplete fragments of otherwise intact phage genomes. Alternatively, some might correspond to defective interfering particles generated during replication (26) or satellites that hijack the RdRp of a helper phage to replicate (27). Distinguishing between these possibilities will be the focus of future work.

### Constructing a ssRNA-phage database

Because our conservative Z ≤ -15 filter captured only about half of the known phages in each metatranscriptome, we reasoned that additional undiscovered phages might still be hidden elsewhere in the data. To recover these additional phages, we performed an exhaustive coat-protein search: we combined the sequences and predicted structures of all newly discovered coats with those of previously identified phages and used them as queries in iterative sequence- and structure-similarity searches across the full Neri and Hou datasets (see **Methods**). We also included previously identified ssRNA phages reported by Edgar et al. (8) Zayed et al. (9), and Chen (28), as well as two databases of well-annotated ssRNA phages from NCBI (29) and IMG/VR v4 (30). Through this process, we identified an additional 83,404 previously unclassified contigs within the Neri and Hou datasets that encoded a properly-folded coat protein, bringing the total number of newly discovered phage contigs to 94,920.

We compiled these results into a database, the San Diego State University Coat Information Bank (“SCIB”), which integrates sequence and structural information for both newly discovered and previously described ssRNA phages (**Supporting Information, Datasets S1 and S2**). For 467,168 distinct phage contigs (94,920 new and 342,248 previously known) and their 100,403 unique coat proteins (52,759 new and 47,644 previously known), we compiled the RNA sequence, predicted RNA secondary structure, and RNA folding stability, as well as the coat-protein sequence, predicted protein structure, and annotations of the encoded coat within the genome. This resource is intended to support basic research in microbial ecology by facilitating the discovery and characterization of new ssRNA phages. It is also intended to support applied research by providing a diverse set of coat proteins with validated folds for the design and production of new capsid-based nanomaterials (**Supporting Information, Fig. S5**).

### Newly discovered ssRNA phages as a source of nanomaterials

To demonstrate the diversity of capsid nanomaterials that can be produced from the SCIB database, we performed a high-throughput protein expression experiment involving 12,000 distinct coats (**Fig. 4A**). We constructed a pET-plasmid library in which each plasmid encodes an isopropyl β-D-1-thiogalactopyranoside (IPTG)-inducible T7 promoter, a unique coat-protein gene, and a 16-nucleotide barcode sequence (**Supporting Information, Fig. S6**). We introduced the plasmid library into E. coli BL21(DE3) by electroporation and induced coat-protein expression by adding IPTG. Following 4 h of expression at 37 °C, we lysed the cells and collected any assembled capsids using established protocols involving multiple rounds of nuclease treatment and density-gradient centrifugation (31).

**Fig. 4.**
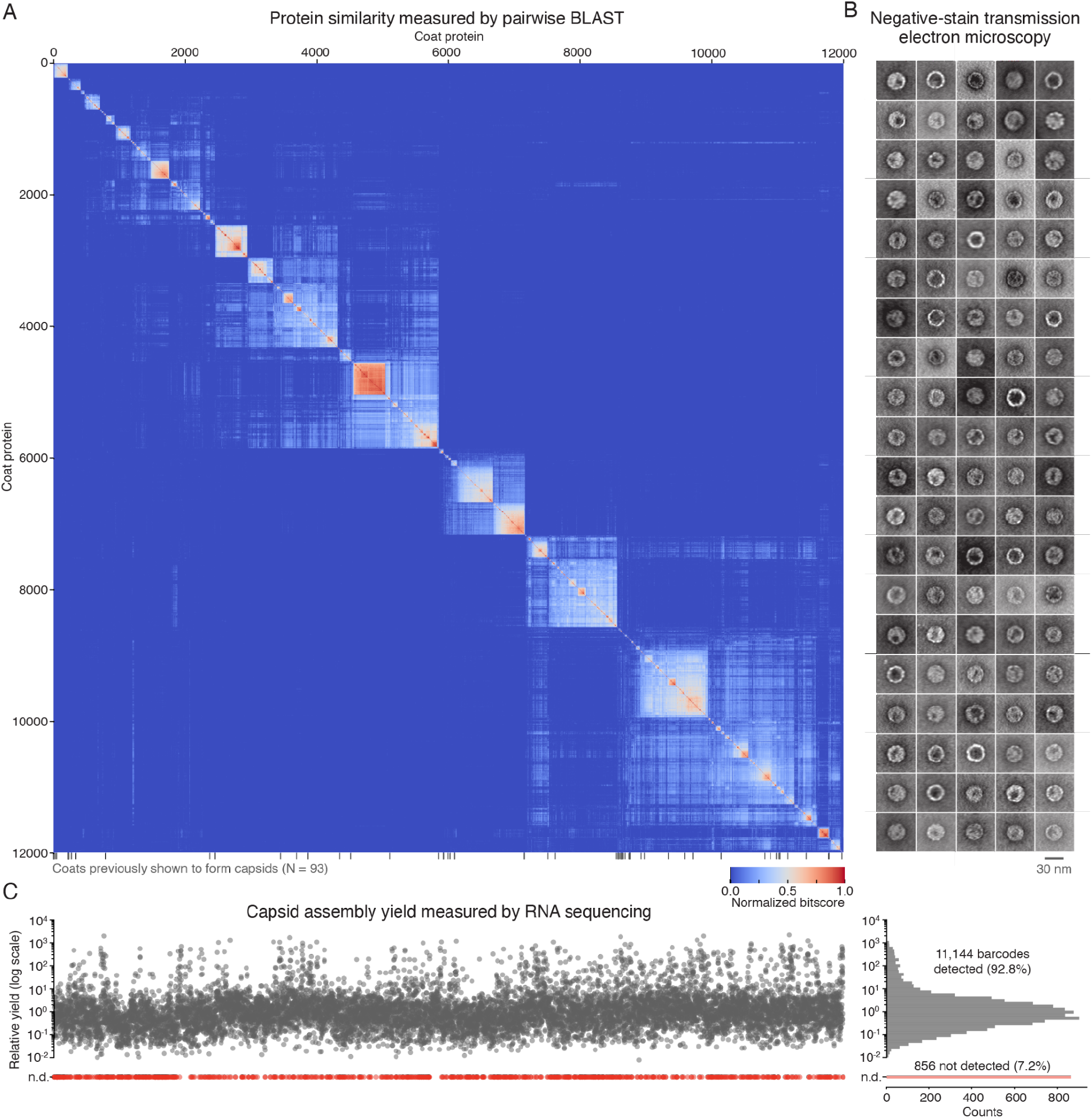
Largescale parallel expression of coat-protein capsids. We expressed a library of 12,000 coat proteins in E. coli and determined capsid assembly yields by sequencing the packaged mRNA. **A**. All-vs-all pairwise BLAST bitscores of the coat library clustered by sequence similarity. Each score was normalized by the maximum self-score of that pair, such that each value lies between 0 and 1. Also included are 93 coats that have been previously shown to form capsids (gray ticks). **B**. Transmission electron microscopy images of 100 randomly selected capsids. **C**. The relative yield of capsid assembly for each coat protein in the library was determined by measuring the prevalence of its mRNA sequence in the population of packaged RNA and dividing by the prevalence of its DNA in the plasmid library. **Left:** The relative assembly yield for each coat protein ordered according to its position in the similarity matrix in panel A. Proteins that did not form capsids in detectable yield (not detected, n.d.) are shown in red. **Right:** Histogram of relative yields.

We verified the formation of ordered capsids by negative-stain transmission electron microscopy (TEM). On the whole, the pool of collected capsids displayed roughly spherical morphologies with diameters of about 30 nm (**Fig. 4B**). Some capsids showed signs of polyhedral faceting consistent with icosahedral symmetry, whereas others exhibited knobby surface features indicative of regularly spaced capsomer protrusions. While orientation effects and uneven staining across the sample precluded rigorous classification of distinct structural types, the images suggest morphological diversity within the population.

We determined the identities of the assembled capsids by RNA sequencing. Because the capsids packaged the mRNA transcripts from which their coat proteins were translated, and because the transcripts carry unique barcodes, we could determine which coat proteins had assembled into capsids by extracting the packaged RNA and detecting the barcodes by high-throughput sequencing. Within the pool of assembled capsids, we detected barcodes for 11,144 of the 12,000 coat proteins in our input library, suggesting 92.8% produced detectable levels of assembly (**Fig. 4C**).

Of the proteins that assembled into capsids, the yield of assembly varied widely. We defined the relative assembly yield as the prevalence of each barcode sequence in the pool of packaged RNA relative to its prevalence in the input plasmid library (see **Methods**). The measured yields spanned five orders of magnitude, with the top 1% of coat proteins generating 400-fold more assembly on average than the median protein (**Fig. 4C, right**). Such large variability is not unexpected, as the library encompasses coat proteins from phages that infect different hosts across diverse environments, some of which differ markedly from the conditions tested here.

Despite this variability, we observed some general trends. On average, coat proteins with more similar sequences produced more comparable assembly yields: for example, pairs of coats whose sequences differed by only one amino acid (Hamming distance, H=1) displayed a median 2.5-fold difference in yield, whereas pairs with H=10 differed by 3.0-fold, and pairs with H=100 differed by 4.5-fold (**Supporting Information, Fig. S7**). That said, certain pairs with H=1 showed 100-fold differences in yield, indicating that single-residue substitutions can have profound effects on assembly. Notably, we found no clear correlation between the nature of such single-residue substitutions—that is, the change in charge, hydropathy, or bulkiness of the substituted side chains—and the change in yield. These observations support the view that capsid assembly is a collective process governed by context-dependent interactions between many subunits, and that differences in assembly yield between such systems are not readily explained by the physicochemical properties of individual residues (32).

To place the relative assembly yields on an absolute scale, we selected SCIB_ID_0055846_0000000 as a representative high-yielding variant and measured capsid production in a single-variant expression experiment. We expressed this variant in isolation under the same growth and induction conditions used in the pooled screen, resolved the assembly products by native agarose gel electrophoresis followed by nucleic acid staining, and compared the resulting band intensities to those obtained for MS2, a well-studied benchmark. In our hands, MS2 yields around 10 mg of capsid per liter of culture under these conditions. SCIB_ID_0055846_0000000 produced bands of comparable intensity to MS2, indicating a similar absolute yield (**Supporting Information, Fig. S8**). Because its relative yield of 129 measured by barcode sequencing places SCIB_ID_0055846_0000000 in the top 5th percentile of the library (**Fig. 4C, right**), we infer that other variants in this upper tail also produced on the order of 10 mg/L capsid, whereas most other variants produced substantially less. Variant-specific optimization of expression and assembly conditions is therefore required to achieve production levels suitable for materials applications across the full library.

### Structural and biophysical characterization of capsids formed from the coat proteins of one newly discovered phage

We expressed the coat protein from a different phage—SCIB_ID_0008658_0000000, which we nicknamed “eliophage”—in isolation to examine its structural and biophysical features in finer detail. Elioiphage coat protein expressed at similar yield to MS2 and assembled into capsids that were readily purified using standard protocols. Negative-stain TEM confirmed the formation of well-ordered capsids (**Fig. 5A**), and cryo-electron microscopy revealed multiple morphologies—including a dominant T = 3 icosahedron, a T = 4 icosahedron, and a rare prolate form (**Fig. 5B**). Consistent particle size and mass distributions measured by TEM and interferometric scattering microscopy (iSCAT) verified the relative abundance of these morphologies within the population (**Figs. 5C and 5D**).

**Fig. 5.**
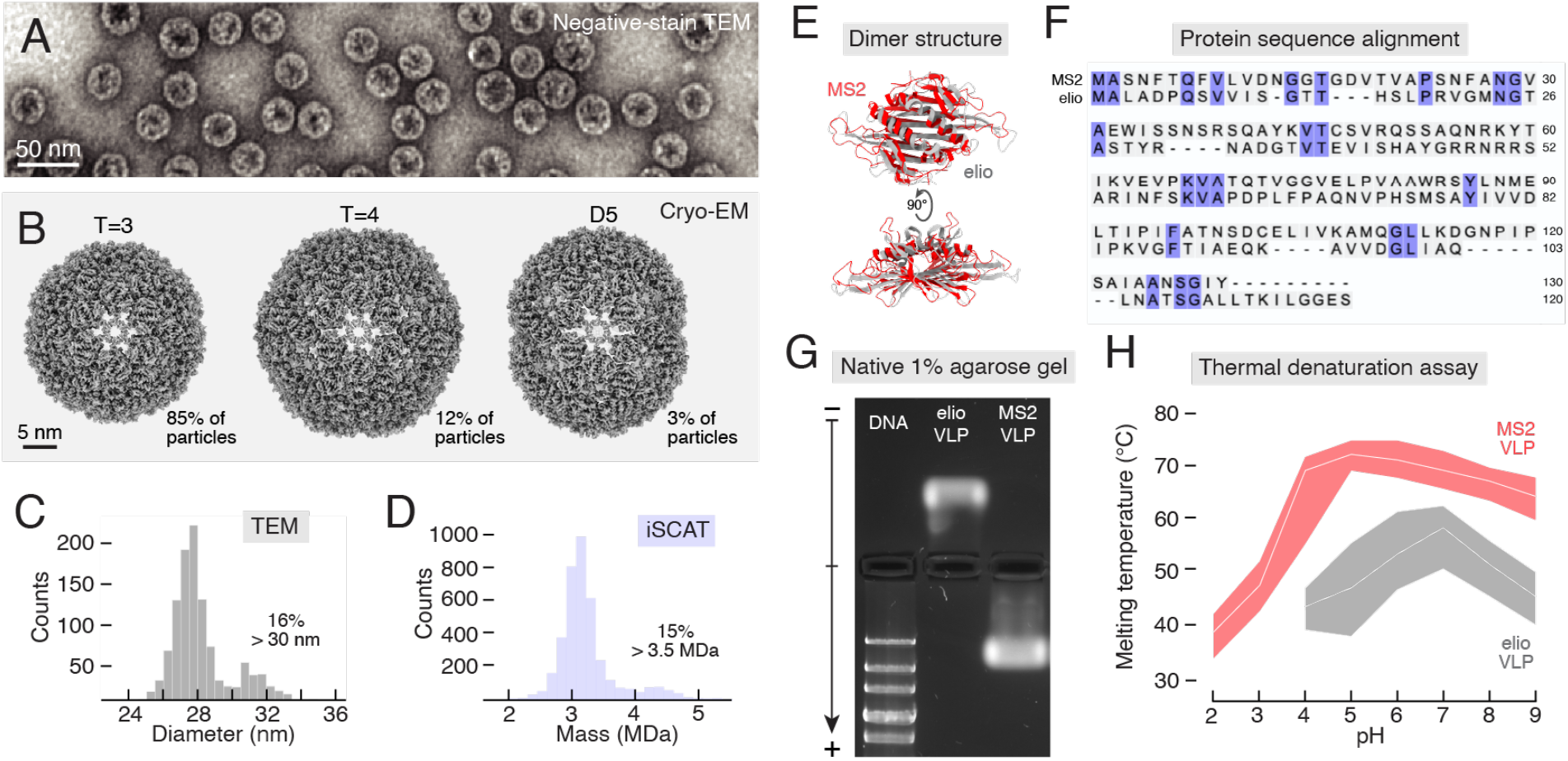
Biophysical characterization of a newly discovered capsid. **A**. Negative-stain TEM of eliophage (SCIB_ID_0008658_0000000) capsids expressed in E. coli. **B**. Cryo-electron microscopy analyzed by single-particle reconstruction reveals several distinct capsid structures, including a dominant T=3 icosahedron, a less common T=4 icosahedron, and a rare prolate cage. **C**. Particle-size histograms obtained by negative-stain TEM confirm the prevalence of larger structures. **D**. Particle-mass histograms show a population of large particles in quantitative agreement with cryo-EM and negative-stain TEM analysis. **E**. The cryo-EM-determined structure of the eliophage dimer (gray) is similar to that of the MS2 coat-protein dimer (red) (pdb_00009n41). **F**. However, the eliophage and MS2 sequences share little similarity. **G**. Native agarose gel electrophoresis performed at pH = 4.8 and stained with ethidium indicate opposite electrostatic charges of eliophage and MS2 capsids. DNA ladder shows 20, 10, 7, 5, and 4 kbp. **H**. Differential scanning fluorimetry shows that eliophage capsids denature at lower temperatures than MS2 capsids do, across all pH values tested. White lines show the peak of the melting transition, and shaded regions show the full-width at half maximum.

Although eliophage and MS2 coat proteins share similar tertiary and quaternary structures (**Fig. 5E**), they differ markedly in sequence and biophysical properties. At the sequence level, eliophage and MS2 coats share only 18% amino-acid identity (**Fig. 5F**). Biophysically, native agarose gel electrophoresis showed that eliophage and MS2 coats carry opposite net surface charges (**Fig. 5G**). Furthermore, thermal stability assays revealed distinct pH-dependent denaturation profiles: eliophage capsids denatured at significantly lower temperatures than MS2 capsids did, across all pH values tested (**Fig. 5H**). These differences point to distinct surface chemistries and inter-subunit interactions—properties that could be relevant to applications in gene delivery because they can influence biodistribution (33), cellular uptake (34), and cargo release (35).

### Newly discovered phage capsids as tools for RNA packaging

To demonstrate that eliophage capsids can package heterologous RNA in vitro, we disassembled the particles, purified the coat proteins, and reassembled the purified protein around foreign RNA. Using MS2 RNA as a model cargo, we confirmed RNA packaging by gel-shift assay and demonstrated nuclease protection of the packaged RNA by survival of the ethidium-stained band following treatment with RNase A (**Fig. 6A**). These results indicate that RNA is both packaged and protected inside of reassembled capsids. TEM analysis further confirmed that the reassembled particles matched the size and curvature of cell-born capsids (**Figs. 6B and 6C**). Together, these experiments show that eliophage coat proteins are not only capable of assembling into stable capsids in cells but can also be directed to package non-native RNAs in vitro, providing a versatile platform for programmable RNA encapsidation and a possible tool for RNA delivery.

**Fig. 6.**
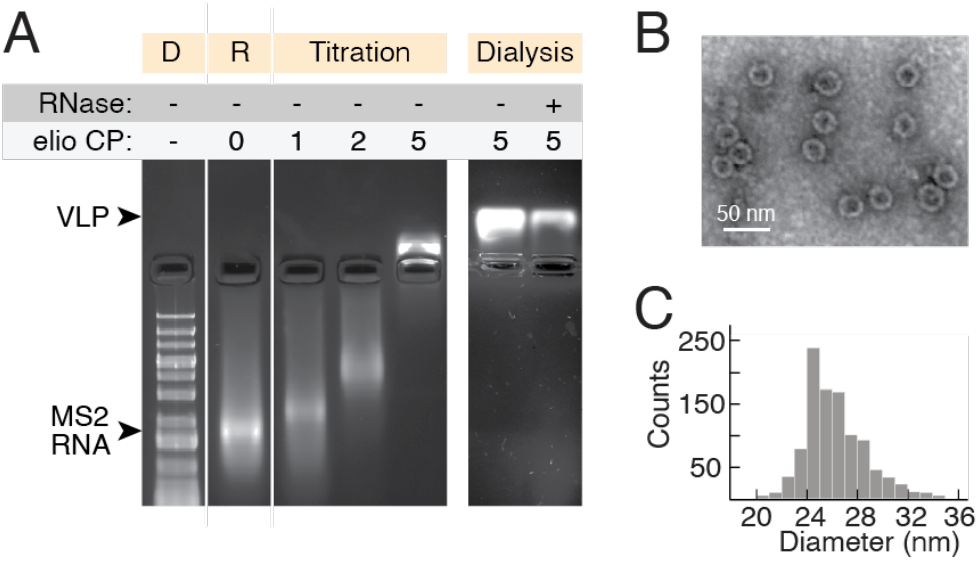
Newly discovered capsids can be disassembled and reassembled around foreign RNA. **A**. Native agarose gel of reassembly of eliophage (SCIB_ID_0008658_0000000) coat proteins (CP) around MS2 RNA. RNA encapsidation was confirmed by gel-shift and resistance to RNase digestion. **B**. Negative-stain TEM images of reassembled capsids show particles with size and curvature comparable to capsids produced in cells. **C**. Size distributions of reassembled capsids measured from TEM images.

## Conclusions

We have shown how RNA-structure prediction can be used to discover new viruses and clarify evolutionary relationships between them. As proof of concept, we used a single, coarse descriptor of RNA structure—MFE Z-score—to detect new ssRNA phages and corroborate their relatedness to narnaviruses, mitoviruses, and botourmiaviruses. Future approaches could incorporate finer structural features—such as specific motifs or higher-order arrangements thereof—to detect and classify other families of viruses and virus-like elements.

Realizing the full potential of RNA-structure-based virus discovery, however, will require better tools for predicting the structures of long RNA molecules, including tertiary and quaternary structures, as has been recently demonstrated for proteins with tools like AlphaFold (36). Developing AlphaFold-caliber tools for RNA structure prediction will require more experimentally determined structures of long RNA molecules for use as training data (37). To date, some of the best experimentally determined structures of long RNAs have come from the packaged genomes of ssRNA phages (38–40). Determining the structures of additional ssRNA phages— including, for example, the yet uncultured phages described here—could help provide the missing data needed to usher in the next generation of RNA structure prediction.

Our results extend those of Liekniņa et al. (10) and Rūmnieks et al. (41), who first showed that coat proteins from uncultured ssRNA phages can assemble into capsids when expressed in E. coli. By measuring capsid assembly yields for thousands of distinct coat proteins, we have generated a quantitative map of the assembly landscape across a vast region of sequence space. The absence of clear features distinguishing high-from low-yielding sequences reinforces the idea that capsid assembly is a non-trivial process whose outcomes are difficult to predict. Future experiments that systematically vary expression conditions, combined with modern tools for high-dimensional data analysis, could lead to models that predict whether a given coat protein will assemble in high yield—a first step toward building a general understanding of the assembly process.

Beyond assembly, our experimental framework opens up new ways of testing capsid functionality. By using packaged RNA barcodes as a readout of capsid prevalence, we envision high-throughput screens in which preassembled capsid libraries are subjected to defined challenges—such as physicochemical stresses or exposure to specific biomolecules or cell types—and survivors are identified by sequencing. Such experiments could enable systematic characterization of particle stability across this rapidly expanding class of nanomaterials. While we have already identified one candidate capsid with favorable traits—including positive surface charge, reduced stability at low pH, and the ability to disassemble and reassemble around foreign RNA—other capsids in our library likely possess additional traits that remain to be identified. The stage is set to turn this remarkable sequence diversity into functional phenotypes.

## Methods

### RNA folding calculations

#### Computing MFE Z-scores

To compute the MFE of an RNA sequence, we used RNAfold from the ViennaRNA software package (2.4.16) (19), with default parameters (RNAfold --noPS sequence_input.fasta > rnafold_output.txt).

To compare MFE values across different RNA sequences, we computed Z-scores. Traditionally, Z-scores are computed by comparing the MFE of a given sequence to the distribution of MFEs obtained from many random shuffles of that sequence, according to the formula:

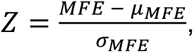

where *μ*_*MFE*_ and *σ*_*MFE*_ denote the mean and standard deviation of the MFEs of shuffled sequences. Different shuffling strategies—such as preserving nucleotide, dinucleotide, or local GC composition—have been shown to affect MFE Z-scores (42–44), but we found these effects are small compared to the magnitude of the Z-scores of ssRNA-phage genomes (**Supporting Information, Fig. S9**). We therefore adopted the simplest possible shuffling strategy: preserving only nucleotide composition.

To accelerate the Z-score calculation, we precomputed MFE distributions for a library of random sequences with lengths from 1,000 to 6,500 nucleotides and G+C/A+U ratios from 0.25 to 1.75, which enabled us to parametrize *μ*_*MFE*_ and *σ*_*MFE*_ as functions of length and nucleotide composition (**Supporting Information, Fig. S10**). Using this precomputed library, we could determine Z-scores for contigs directly from their MFEs, without generating and folding new shuffled sequences.

#### Comparing MFE Z-scores across known RNA virus families

To compare MFE Z-scores across different families of RNA viruses, we used the Riboviria database (riboviria.org) (7). For each family in the database, we randomly selected up to 100 complete genome sequences, computed their MFE values, and calculated Z-scores as described above. Families with fewer than three sequences (N < 3) were excluded. To maintain consistency with our precomputed parameters, genomes longer than 6,500 nucleotides were trimmed to this length prior to analysis.

#### Computing MFE Z-scores across metatranscriptomic datasets

We computed MFE Z-scores for two published metatranscriptomic datasets: a 5.1×10^6^-contig dataset reported by Neri et al. (7), and a 7.1×10^8^-contig dataset reported by Hou et al. (12). To facilitate Z-score calculations, we excluded contigs containing noncanonical nucleotides and restricted contig lengths to 1,000-6,500 nucleotides and G+C/A+U ratios to 0.25-1.75. This left 4.7×10^6^ contigs in the Neri dataset and 1.2×10^8^ contigs in the Hou dataset (**Supporting Information, Fig. S11**). These datasets contained both the contig sequences and their RdRp-based taxonomic annotations.

### Identifying and cataloging ssRNA-phage coat proteins

#### Identifying coat proteins

To identify coat proteins, we performed multiple rounds of sequence-similarity detection and structural validation, as detailed in **Supporting Information**. Briefly, to detect sequence similarity, we used a combination of tBLASTn alignment (45), MMseqs2 easy-search (v15-6f452) (46), and DIAMOND clustering (v2.1.9) (47) against validated coats. Clustering was performed iteratively from 90% to 10% similarity in 10% increments, following the approach of Hou et al. (12). Protein-coding regions were predicted using Prodigal (v2.6.3) (48). Structural validation was performed using AlphaFold2 (49). Structures were predicted using ColabFold (v1.5.5) (50), with all proteins treated as homodimers. Multiple sequence alignments were custom built from validated coat-protein sequences using Clustal Omega (v1.2.4) (51) and formatted as .a3M files using HHsuite (v3.3.0) (52). Predicted structures were aligned to validated coats using Foldseek (v8.ef4e960) (53): proteins with TM-scores > 0.8 were automatically retained; those scoring between 0.4 and 0.8 were manually inspected for evidence of the canonical fold; and those scoring below 0.4 were discarded.

#### Building a coat-protein database

To build our SCIB database, we screened multiple metatranscriptomic datasets using the coat-protein identification strategy described above. In addition to the Neri and Hou datasets, we screened collections of previously identified ssRNA phages reported by Edgar et al. (8) Zayed et al. (9), and Chen et al. (28), as well as well-annotated ssRNA phages from the NCBI Virus database (accessed February 2, 2024) (29) and the IMG/VR v4 database (accessed June 11, 2025) (30). Through this process we identified 100,403 unique coat proteins. For each, we recorded the protein sequence and its predicted structure, along with sequence and structural information about the contig from which it was derived, including the RNA sequence, predicted secondary structure, MFE, and Z-score MFE. In addition, we also included previously identified contigs derived from ssRNA phages from the above sources. Each contig in the SCIB database received a unique SCIB identification number, SCIB_ID_XXXXXXX_YYYYYYY, in which XXXXXXX designates a unique coat protein sequence and YYYYYYY designates a unique contig sequence (**Supporting Information, Datasets S1 and S2**).

#### Constructing a plasmid library for capsid assembly experiments

We selected a subset of 12,000 coat-protein variants from our SCIB database for expression in E. coli. Proteins were randomly sampled from the database and retained if they met the following criteria:

1. Length between 119 and 143 amino acids. Shorter proteins were discarded to avoid incomplete sequences; longer proteins were incompatible with DNA synthesis.
2. No cysteine residues. Proteins containing one or more cysteines were discarded to avoid spurious disulfide bond formation.
3. High-confidence structural predictions. Proteins with low pLDDT scores at the N-terminus were excluded to reduce errors stemming from incorrect start codon prediction.
4. Canonical secondary structures. Proteins were required to match the secondary structure pattern of the MS2 coat protein: six N-terminal β-strands followed by two C-terminal α-helices. Sequences predicted to contain additional structural elements were excluded to avoid atypical folds that might express poorly. Secondary structure annotations were derived from AlphaFold2-predicted models using the DSSP module (54) in Biopython (55).

A pET plasmid library (Twist Bioscience) was constructed encoding 12,000 retained proteins. In addition to a unique coat-protein gene, each plasmid insert included a unique 3’ barcode sequence (**Supporting Information, Fig. S6**).

### Largescale capsid assembly experiments

#### Protein expression

We introduced the plasmid library into E. coli BL21(DE3) cells by electroporation: 1 µg of plasmid DNA was mixed with 40 µL of electrocompetent cells (Intact Genomics), transferred to a pre-chilled 0.1-cm gap electroporation cuvette (Bio-Rad), and electroporated using the EC1 preset program (1.8 kV, exponential decay pulse) on a MicroPulser electroporation system (Bio-Rad). We performed three electroporations and pooled the transformed cells. We determined the number of transformants to be greater than 10^8^ by plating serial dilutions on selective media and counting colony-forming units. The transformed cells were added to 2L of LB media containing 0.05 mg/mL kanamycin and incubated at 37°C with shaking at 250 rpm. When the optical density at 600 nm reached 0.5, we removed 1L of the culture for plasmid extraction using a Plasmid Maxi Kit (QIAGEN), following the manufacturer’s instructions. The remaining 1L of culture was induced with 1 mM of IPTG and incubated for 4 h at 37°C with shaking at 250 rpm. The cells were pelleted, resuspended in PBS buffer (10 mM phosphate, pH=7.4; 137 mM NaCl, and 3 mM KCl) (Fisher), and mechanically lysed by sonication.

#### Capsid purification

Capsids were purified from cell the lysate using previously established techniques (31). All steps were performed in PBS buffer, except steps involving nuclease treatment, which were performed in PBS supplemented with 1× DNase-I-buffer (NEB). We performed nuclease treatment of the lysate using 5 μg/mL of RNase A and 40 U/mL DNase I for 1 h at 37°C and then overnight at room temperature. Sucrose density centrifugation was used to separate the nuclease-resistant capsids and their packaged RNA from the digested material: we layered 12 mL of treated sample on 1 mL of 15% (w/v) sucrose solution in Ultra-Clear centrifuge tubes (Beckman Coulter) and spun the tubes at 36,000 rpm for 6 hours at 4°C in a SW 41 rotor (Beckman Coulter). The supernatant was decanted and the resuspended pellets were subjected to 3 rounds of dialysis against fresh PBS buffer using the 3.5-kDa dialysis cups (Thermo Scientific). Next, we performed another nuclease treatment using 5 μg/mL of RNase A and 40 U/mL DNase I for 1 h at 37°C. Here, a 0.5-mL 100-kDa centrifugal filter unit (MilliporeSigma) was used to remove the digested material: five rounds of filtration were performed in which the sample volume was increased to 0.5 mL with fresh PBS buffer and then reduced by at least 5-fold by spinning at 14,000 rpm for 5 min at room temperature. Finally, we performed chloroform extraction to remove lipids.

#### RNA extraction

Packaged RNA was extracted from purified capsids using an RNeasy Mini Kit (QIAGEN), following the manufacturer’s instructions. RNA integrity was determined by capillary gel electrophoresis (Agilent Tapestation), and purity with respect to protein contamination was determined by UV spectro-photometry. Extracted RNA was stored in TE buffer (10 mM Tris, pH 7.0; 1 mM EDTA) at -80 °C prior to sequencing.

#### Sequencing

We used high-throughput short-read sequencing to quantify the prevalence of each barcode sequence in (i) the plasmid DNA purified from transformed cells harvested immediately prior to IPTG induction, and (ii) the RNA extracted from purified, nuclease-treated capsids. Library preparation and 150-bp paired-end sequencing were performed by Azenta Life Sciences, and data were returned as gzipped FASTQ files. We deposited these files into the Sequence Read Archive (SRA) (BioProject number: PRJNAXXXXX [will deposit upon publication]).

Our analysis of the sequencing data is detailed in **Supporting information**. Briefly, we compiled a FASTA file containing the 32-nt terminal window of each coat-protein insert— comprising the last 16 nt of the coat gene and the 16-nt barcode—and used bowtie2-build (v2.4.5) (56) to create the index for alignment. Reads were quality-filtered using fastp (v0.23.4) (57) with default parameters. We aligned each read separately, meaning we did not make use of the paired-end nature of the data. Alignment was performed using Bowtie2 in local alignment mode with strict seed parameters (--local --very-sensitive-local -N 0 -L 32 -k 1). Resulting SAM files were converted to coordinate-sorted BAM files using samtools (v1.16.1) (58) view and samtools sort. Mapped-read counts for each barcode were obtained from the sorted BAM files using samtools idxstats. This procedure was performed separately for plasmid-DNA libraries and packaged-RNA libraries (**Supporting Information, Fig. S12**).

#### Assembly yield calculation

We defined the assembly yield for each coat-protein variant, *i*, as:

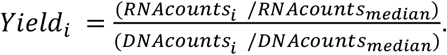

Here, *RNAcounts*_*i*_ is the mapped-read count of barcode *i* in the RNA library, and *RNAcounts*_*median*_ is the median RNA count across all barcodes. Likewise, *DNAcounts*_*i*_ is the read count of barcode *i* in the DNA library, and *DNAcounts*_*median*_ is the median DNA read count. Normalizing by the median corrects for differences in sequencing depth between samples while reducing the impact of high-count outliers. The resulting yield provides a relative measure of how efficiently each variant packages its own RNA transcript into a nuclease-resistant capsid. Variants with zero DNA counts (2 variants out of 12,000) were excluded from the analysis.

### Biophysical characterization of capsids

#### Negative-stained transmission electron microscopy (TEM)

We performed TEM imaging as described previously (31). Briefly, a 6-µL drop of approximately 10 nM particles in either PBS or TNE buffer (50 mM Tris, pH 7.0; 100 mM NaCl; 1 mM EDTA) was placed on Parafilm (Bemis), and a freshly glow-discharged carbon-coated 200-mesh copper grid (Electron Microscopy Sciences) was floated on the drop, carbon side down, for 2 min. The grid was removed, blotted with filter paper, and placed onto a 10-µL drop of 1% (w/v) uranyl acetate for 30 s, blotted, and transferred onto a fresh 1% uranyl acetate drop for an additional 30 s. After blotting, we imaged the grids using a Tecnai 12 transmission electron microscope (FEI) operating at 120 kV and equipped with a side-mount AMT camera. Images were recorded at 97,000× magnification and analyzed using FIJI (59) to determine particle size distributions.

#### Cryo electron microscopy (cryo-EM)

We applied 5-µL aliquots of purified capsids to glow discharged grids (either Quantifoil R 1.2/1.3 or Quantifoil R 1.2/1.3 coated with a layer of ultra-thin carbon). The grids were then plunge frozen in liquid ethane using a Vitrobot Mark IV operated at 4 °C and 100% humidity, with a blot force of 1 and 5 s of blotting time. We collected cryo-EM data using a Talos Arctica (Thermo-Fisher) operating at 200 keV with Selectris energy filter (slit width: 10 eV) and equipped with a Falcon 4i direct electron detector. 5,279 micrographs were collected at 130,000× magnification (0.886 Å/pixel) by recording EER frames over 6.00 s for a total dose of 44.4 e−/Å^2^. Data processing was carried out using Cryosparc v4.4.1 (60). Dose-fractionated movies were subjected to motion correction and CTF estimation was performed on the resulting images. Particles were picked using blob picking followed by template picking. 287,912 particles were used for 3D refinement for T=3 capsids, 48,245 particles were used for T=4 capsids, and 10,130 particles were used for D5 capsids. The overall map resolutions were estimated based on the gold-standard Fourier shell correlation (FSC 0.143) (61). The T=3 icosahedral map was determined to a resolution of 2.4 Å, the T=4 icosahedral map was determined to 2.8 Å, and the D5-symmetric map to 3.4 Å (**Supporting Information, Table S1**). These maps were deposited into the Electron Microscopy Data Bank (EMD-XXXXX, EMD-XXXXX, EMD-XXXXX [will deposit upon publication]). We generated initial models of the icosahedral maps using ModelAngelo (62), using both sequence and nonsequence modes. Refinement was carried out using Phenix (63) and model adjustments were made in COOT (64). Models were deposited into the Protein Data Bank (PDB ID: XXXX, and XXXX [will deposit upon publication]).

#### Interferometric scattering microscopy (iSCAT)

We determined the mass distribution of the purified capsids using iSCAT microscopy as described previously (31). Briefly, capsids were diluted in water and added to a glass coverslip on a TwoMP microscope (Refeyn). Individual particle-coverslip binding events were detected as changes in image contrast. To convert the contrast signal into absolute mass, we generated a calibration curve using standards of known molecular weight. We used jack bean urease (Sigma-Aldrich), which assembles into defined oligomeric species with discrete masses (270, 540, 820, and 1,080 kDa). In addition, we included virus-based standards consisting of empty MS2 coat-protein capsids (2,470 kDa) and intact MS2 virions (3,600 kDa). These reference measurements produced a linear scaling between iSCAT contrast and particle mass, which we applied to determine the masses of purified capsids.

#### Differential scanning fluorimetry (DSF)

We used DSF to measure the thermal stability of capsids across a range of pH values, as described previously (65). DSF monitors the fluorescence of SYPRO Orange dye (Molecular Probes), which brightens upon binding to hydrophobic regions of coat proteins that become exposed during temperature-induced capsid denaturation (66). MS2 and eliophage capsids were diluted to 0.025 mg/mL in UB4 buffer (35 mM HEPES, 35 mM MES, and 35 mM sodium acetate) (67) prepared at a range of pH values between 2 and 9. We added SYPRO Orange to a final concentration of 2.5×. Samples were loaded into a CFX Connect real-time PCR instrument (Bio-Rad) and subjected to thermal scanning from 25 °C to 95 °C in 1 °C increments, with a 30-second equilibration period at each step. Fluorescence was monitored at each temperature using excitation/emission wavelengths of 470/550 nm. We performed experiments in quadruplicate, and averaged the fluorescence traces across replicates. For each condition, we defined the melting temperature as the temperature corresponding to the maximum of the first derivative of the fluorescence signal. The full width at half maximum of the peak was used to characterize the temperature range over which disassembly occurred.

### RNA packaging

#### Reassembly of eliophage capsids around foreign RNA in vitro

In-vitro reassembly was performed by mixing purified eliophage coat protein with heterologous RNA. We prepared RNA by in vitro transcription and purified it using the RNeasy Mini Kit (QIAGEN), following the manufacturer’s instructions. Eliophage coat protein was prepared by treating cell-purified eliophage capsids with ice-cold glacial acetic acid and removing the RNA component by centrifugation, as described previously (68). We measured eliophage protein concentration using a Bradford assay (Pierce) and determined the extinction coefficient at 280 nm of the coat-protein dimer to be 9,100 M^-1^ cm^-1^. Purified protein was stored in 20 mM acetic acid at 4 °C and discarded after 2 weeks.

For initial assembly experiments, eliophage coat protein and RNA were mixed in TNE buffer. The protein concentration was varied from 1 to 5 μM of dimers while the RNA concentration was held constant at 20 nM. We left the mixtures on ice for 45 min and then analyzed the products by native 1% agarose gel electrophoresis performed in sodium acetate buffer (0.1 M sodium acetate, 1 mM EDTA, pH 4.8). The gel was run at 50 V for 2 h at 4°C, stained with ethidium, and imaged using a ChemiDoc MP imaging system (Bio-Rad). The mobility of the RNA band shifted with increasing protein concentration, but did not reach a position consistent with proper eliophage capsids.

To promote proper capsid assembly, we used dialysis to slowly change the assembly buffer from high-salt TNE buffer (50 mM Tris, pH 7.0; 1000 mM NaCl; 1 mM EDTA), in which protein-RNA interactions are screened, to regular TNE buffer, in which the interactions are turned on. Following 2 h of dialysis at 4°C, the reassembly products co-migrated with proper capsids when analyzed by native 1% agarose gel electrophoresis, and the RNA band was resistant to treatment with RNase A. TEM confirmed the presence of well-formed capsids.

## Acknowledgements

We chose the nickname “eliophage” for SCIB_ID_0008658_0000000 in appreciation of our friend and colleague Moselio “Elio” Schaechter, whose *Small Things Considered* blog (https://smallthingsconsidered.blog/) inspired some of this work. We thank Ben Rogers, Tom Huxford, Elena Rivas, Sean Eddy, and Eugene Koonin for helpful discussions. Some of the research was supported by the National Institute of General Medical Sciences of the National Institutes of Health (R35GM140803 to K.N.P. and R35GM157105 to R.F.G.) and the National Science Foundation (CAREER award number 2443955 to R.F.G.). R.F.G acknowledges support from Ionis Pharmaceuticals. We thank the California Metabolic Research Foundation for its support of biochemical research at San Diego State University.

