## Supporting Information for "Screening metatranscriptomes for ultrastable RNA secondary structures reveals hidden bacteriophages and novel capsid nanomaterials"

**This PDF file includes:**

Supporting text
Figures S1 to S12
Table S1
SI References

**Supporting Text**

**Methods**

**Identifying coat proteins.** To identify coat proteins within metatranscriptomic contigs and previously identified ssRNA-phage genomes, we performed the following steps: **1.** we searched for proteins with sequence similarity to verified coats, and **2.** we predicted the 3D structure of candidate coat proteins to test if they adopt the canonical fold.

**1. Sequence-similarity search.** We used three complementary approaches to identify proteins with sequence similarity to verified coats:

**tBLASTn alignment:**

i. Align the contig pool against validated coat proteins using a tBLASTn search (1), keeping only those contigs with a significant hit ($e < 10^{-3}$).

ii. Predict the full-length protein sequence of each hit using Prodigal (v2.6.3) (2), run in metagenomic mode.

**MMSeqs2 alignment:**

i. Align the contig pool against validated coat proteins using MMseqs2 easy-search (v15-6f452) (3), keeping only those contigs with a significant hit ($e < 10^{-3}$).

ii. Predict the full-length protein sequence of each hit using Prodigal.

**DIAMOND clustering:**

i. Predict the encoded proteins using Prodigal.

ii. Cluster the amino acid sequences of the encoded proteins with those of validated coats using DIAMOND (4), keeping only those proteins that cluster with at least one validated coat. We performed the clustering iteratively, from 90% to 10% similarity, in increments of 10%, as described by Hou et at. (5).

**2. Structure validation.** To test if a protein adopts the canonical fold, we predicted its 3D structure using AlphaFold2 (6) and aligned the structure against those of verified coats using Foldseek (v8.ef4e960) (7), keeping only those proteins with high alignment scores:

i. For structure prediction, we used ColabFold (v1.5.5) (8) with custom MSAs involving all putative coat-protein sequences prepared using Clustal Omega (v1.2.4) (9) and formatted as .a3M files using HHsuite (v3.3.0) (10). Proteins were folded as homodimers, generating 1 model per dimer.

ii. For structural alignments, proteins with TM-scores above 0.8 were automatically kept, proteins with TM-scores of 0.4-0.8 were manually inspected for evidence of the canonical fold, and proteins with TM-scores below 0.4 were discarded.

We screened multiple datasets using the coat-protein identification strategy described above. In addition to the Neri (11) and Hou (5) datasets, we screened collections of previously identified ssRNA phages reported by Edgar et al. (12), Zayed et al. (13), and Chen et al. (14), as well as well-annotated ssRNA phages from the NCBI Virus database (accessed February 2, 2024) (15) and the IMG/VR v4 database (accessed June 11, 2025) (16). Specifically, we applied the protocols described above, iteratively, to different portions of the datasets, as follows:

I. Starting with $4.4\times10^5$ previously identified ssRNA phages within the Neri, Hou, Edgar, and Zayed datasets, and using 1,684 well-annotated ssRNA-phage coat proteins from NCBI as verified coats, we performed DIAMOND clustering and MMseqs2 alignment, followed by structural validation, and identified 132,657 unique contigs encoding 37,810 distinct coats.

II. We then turned to the $2.7\times10^7$ unclassified contigs within the Neri and Hou datasets with lengths of 1,000-6,500 nucleotides and GC/AU ratios of 0.25-1.75. Using the coats identified in the previous steps as verified coats, we performed DIAMOND clustering and MMseqs2 alignment, followed by structural validation and identified 52,457 additional contigs encoding 18,500 additional coats.

III. Next we examined the $1.4\times10^8$ unclassified contigs within the Neri and Hou datasets with lengths of 500-1,000 nucleotides and GC/AU ratios of 0.25-1.75. Using the coats identified in the previous steps as verified coats, we performed DIAMOND clustering and MMseqs2 alignment, followed by structural validation and identified 18,278 additional contigs encoding 18,092 additional coats.

IV. Then we analyzed $9.4\times10^4$ contigs from the IMG/VR v4 database. Using the coats identified in the previous steps as verified coats, we performed DIAMOND clustering and MMseqs2 alignment, followed by structural validation and identified 11,753 additional contigs encoding 6,829 additional coats.

V. Finally, we reexamined the full Neri and Hou datasets against all previously verified coats using tBLASTn alignment followed by structural validation and identified 34,240 additional contigs and 23,925 additional coats.

**RNAseq data analysis.** To quantify the prevalence of each RNA or DNA barcode in the short-read Illumina sequencing data, we performed the following steps: **1.** we generated a reference index based on the terminal 32-nucleotides of each plasmid variant insert, which contains the unique 16-nt barcode sequence; **2.** we filtered the sequencing reads based on quality; and **3.** we aligned the reads against the index to quantify the prevalence of each variant. We performed the same analysis for both RNA and DNA sequencing data.

1. **Generating a reference index.** We compiled the 32-nt 5′-terminal sequences of each insert variant in the plasmid library into a FASTA file and used samtools (v1.16.1) (17) and bowtie2 (v2.4.5) (18) to generate index files for aligning sequencing reads to specific variants:

```
# Generate FASTA index
samtools faidx inserts_term32.fasta
```

```
# Build Bowtie2 index files from the FASTA
bowtie2-build inserts_term32.fasta inserts_term32
```

2. **Read filtering.** We filtered the raw FASTQ files using fastp (v0.23.2) (19) to remove low-quality reads and adapter sequences. Filtering parameters included a minimum read length of 15 bp (-l 15), sliding window size of 8 (-w 8), base quality threshold of 15 (-q 15), maximum of 40% unqualified bases per read (-u 40), no more than 5 ambiguous bases (-n 5), and trimming of a specific adapter sequence (-a CTGTCTCTTATACACATCT):

```
# Filter unpaired reads for overall quality
fastp \
  -i raw_reads_R1.fastq.gz \
  -o filtered_reads_R1.fastq.gz \
  -l 15 -w 8 -q 15 -u 40 -n 5 \
  -a CTGTCTCTTATACACATCT
```

3. **Read alignment and quantification.** Filtered reads were aligned against the index of 32-nt terminal sequences using bowtie2 in local alignment mode (--very-sensitive-local), with a seed length of 32 (-L 32), zero mismatches (-N 0), and a maximum of one alignment reported per read (-k 1). Reads were aligned independently, without using paired-end information.

```
# Filter unpaired reads for overall quality
bowtie2 \
  --local --very-sensitive-local -N 0 -L 32 -k 1 \
  -x inserts_term32 \
  -U filtered_reads_R1.fastq.gz \
  | samtools view -b -F 4 - \
  | samtools sort -o aligned_R1.sorted.bam
```

BAM files were indexed using samtools index and quantified using samtools idxstats:

```
# Index the BAM file and count reads mapping to each insert
samtools index aligned_R1.sorted.bam
samtools idxstats aligned_R1.sorted.bam >
coverage_table.txt
```

The resulting mapped-read counts were used to determine the assembly yield for each variant in the library (see **Main Text**).

### Supporting Figures and Tables

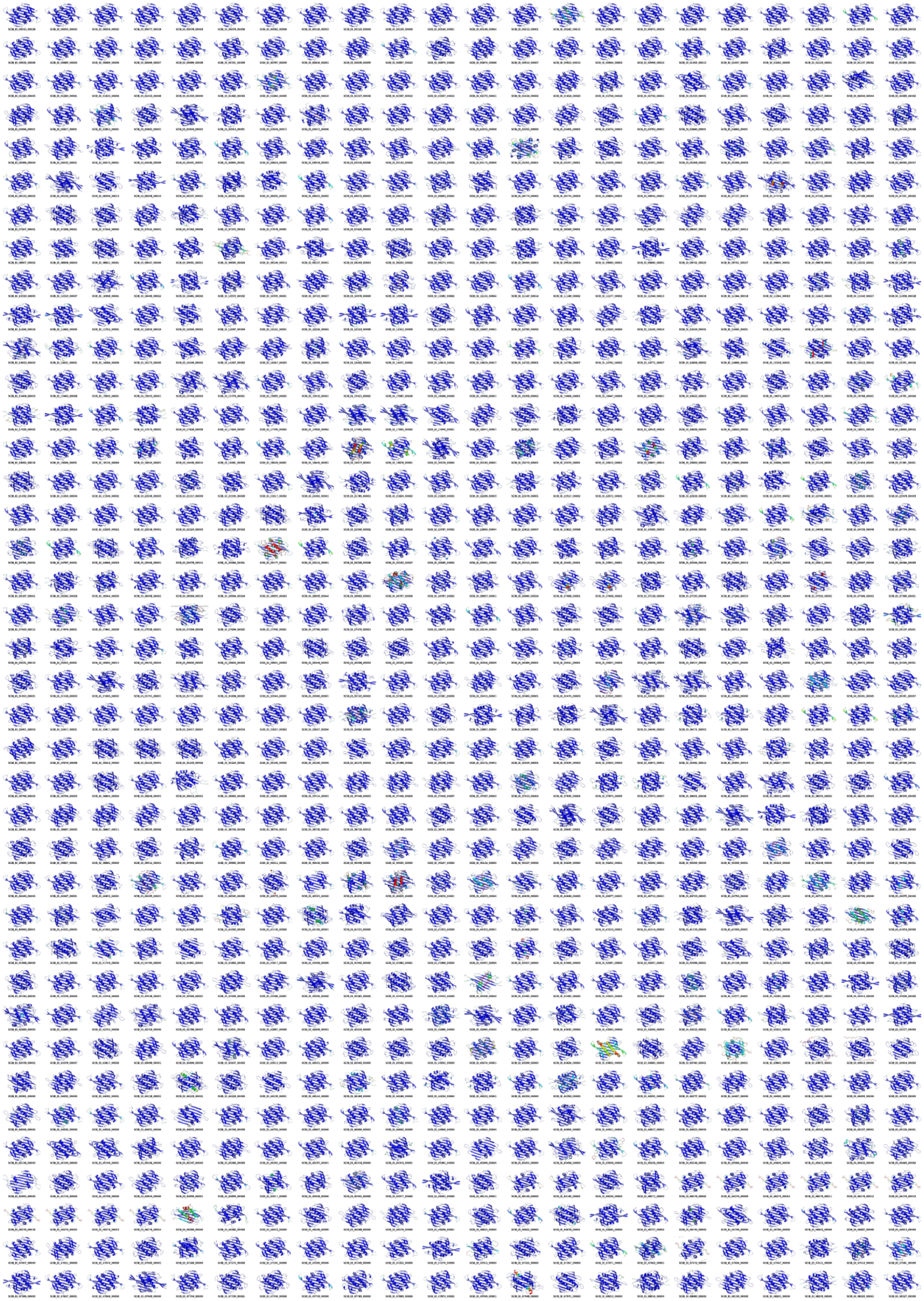

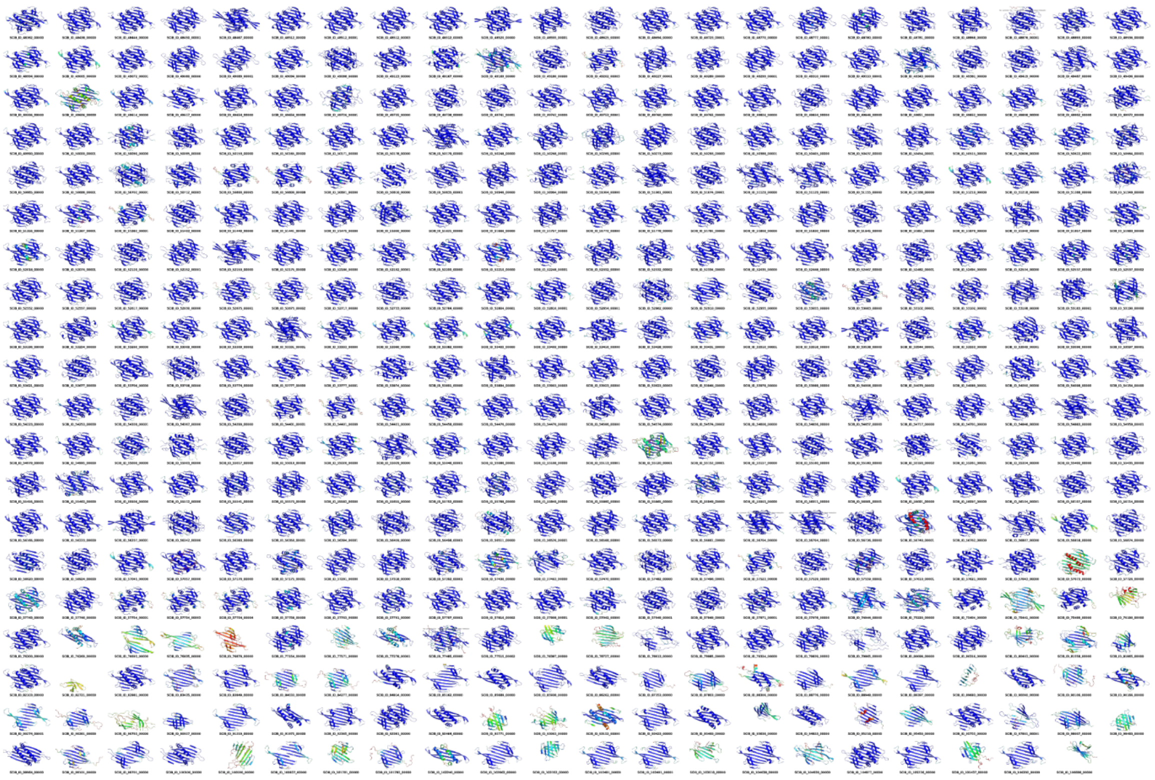

**Fig. S1. AlphaFold2-predicted structures of coat-protein homodimers from previously unclassified ssRNA phages in the Neri dataset.** Structural models are colored by prediction confidence (blue = high confidence, red = low confidence). Associated SCIB ID numbers are displayed below each model.

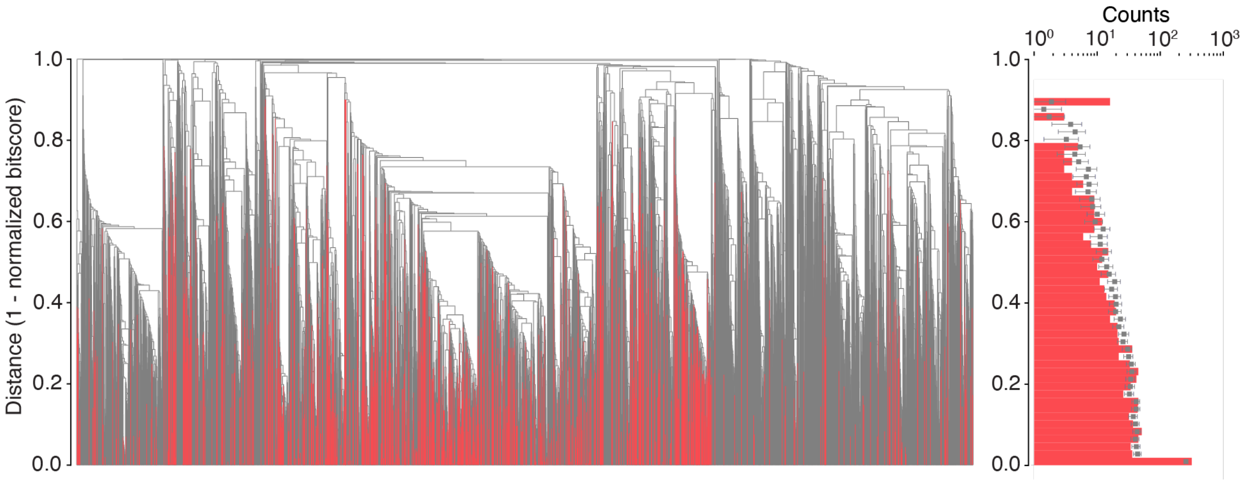


**Fig. S2. Sequence similarity of ssRNA-phage coat proteins in the Neri dataset. Left:** BLASTp dendrogram containing 1,200 newly discovered and 10,712 previously identified coats. We defined distance values as 1-bitscore/self-bitscore and constructed the dendrogram by clustering the distances using the UPGMA algorithm. Newly discovered coats are highlighted in red. The height of each red line reflects the distance at which a newly discovered coat diverges from a node with a previously identified one, such that longer lines indicate coats with increasingly novel sequences. **Right:** Histogram showing the height distribution of red lines in panel A. Gray points and bars show the mean and standard deviation of bin heights determined by randomly sampling 1,200 previously identified coats and determining their divergence from the remaining known coats, as was done in panel A. The distribution obtained for the newly discovered coats is similar to those of randomly sampled known coats, showing that the new coats are just as diverse as the known ones.

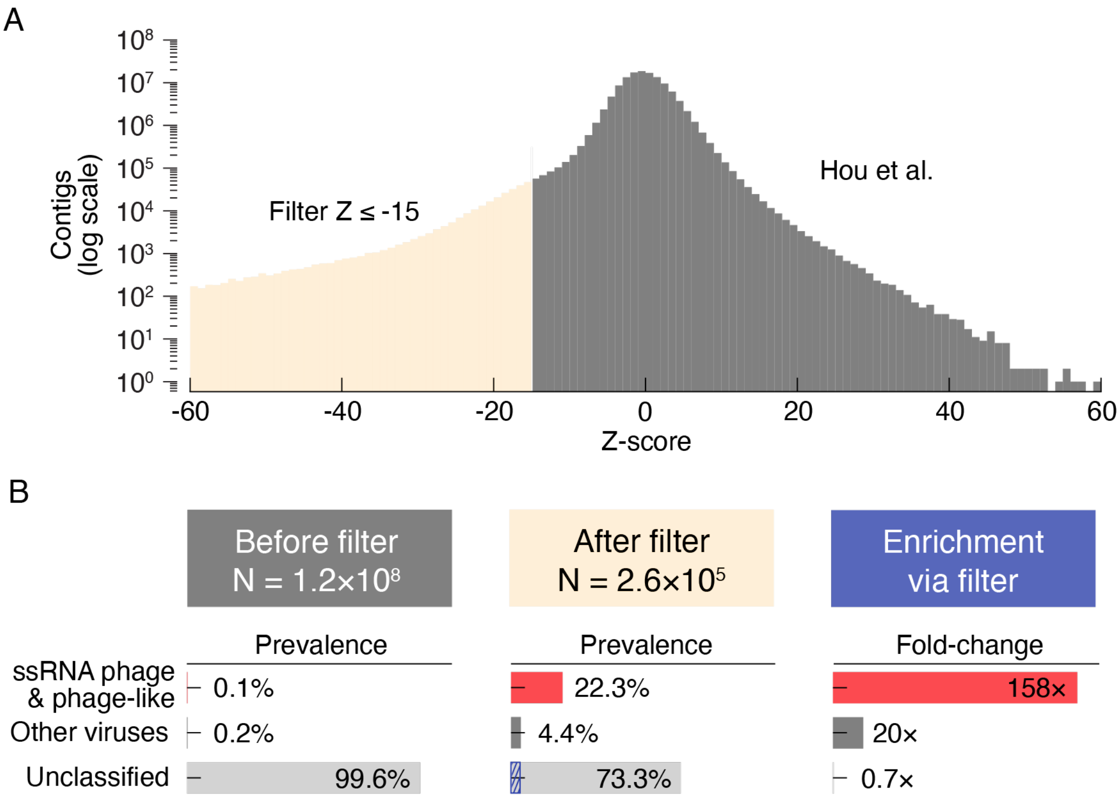


**Fig. S3. Z-score filtering of the Hou dataset reveals known and novel ssRNA-phages. A.** Z-score distribution of the Hou dataset. We filtered the dataset, retaining contigs with Z ≤ -15 (light orange). **B.** The prevalence of ssRNA phages and phage-like viruses (red), other virus families (dark gray), and unclassified contigs (light gray) are shown before and after filtering. Note that phage and phage-like viruses are grouped together, since Hou et al. did not distinguish between these families in their previous analysis. A total of 10,219 previously unclassified contigs retained by the filter were found to encode proteins with the canonical ssRNA-phage coat-protein fold (blue stripes).

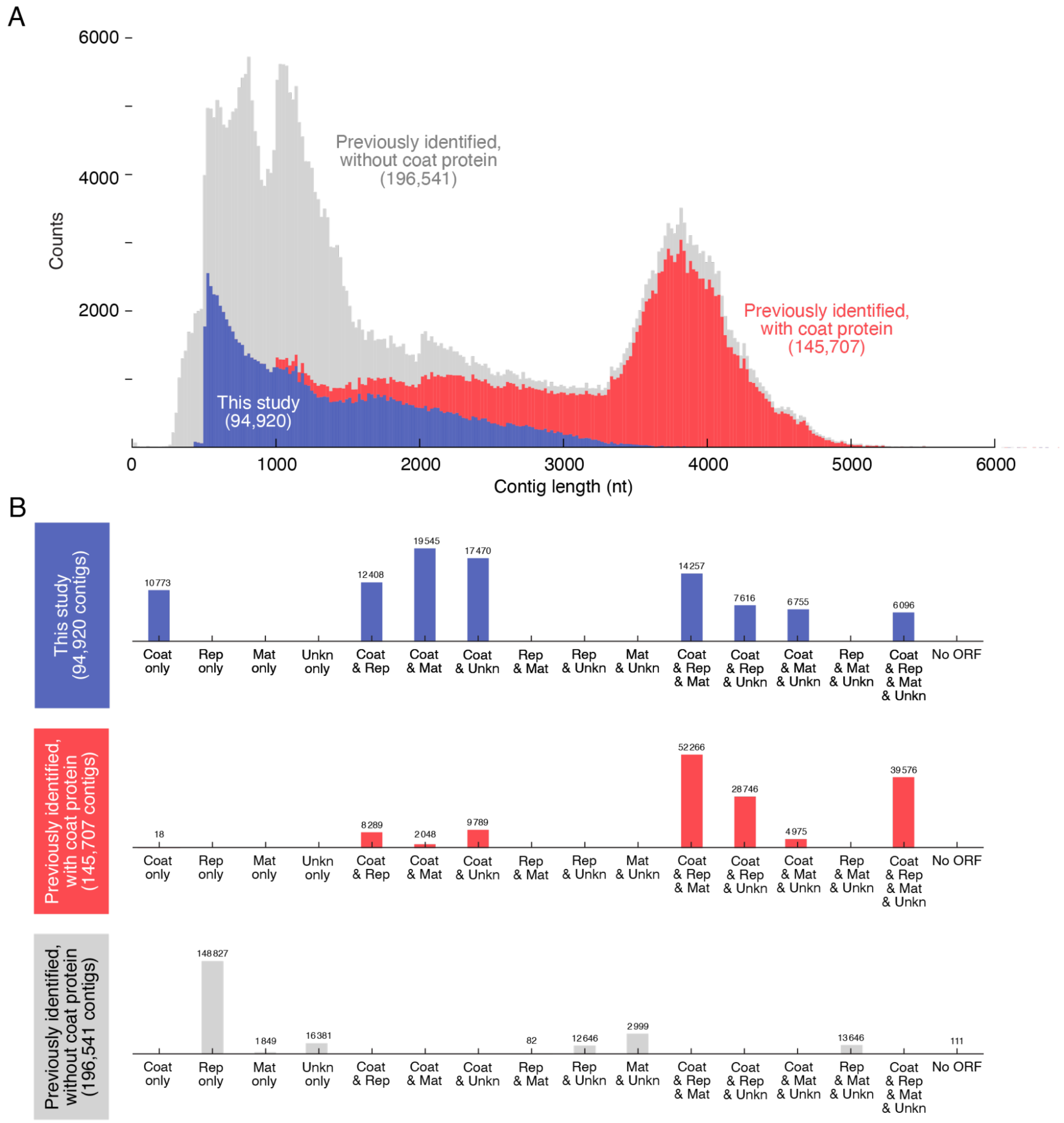


**Fig. S4. Contig length and gene content of newly discovered and previously identified ssRNA phages**. **A.** Stacked histogram of contig lengths: blue bars show new ssRNA-phage contigs identified in this study; red bars show contigs previously identified as ssRNA phages that contain coat genes; and gray bars show contigs previously classified as ssRNA phages that lack coat protein genes. Open reading frames were predicted using Prodigal and annotated based on sequence similarity to known phage proteins. **B.** Bar plots show the number of contigs containing different combinations of Coat, RdRp (Rep), maturation (Mat), and unknown (Unkn) proteins. **Top:** new ssRNA-phage contigs identified in this study. **Middle:** contigs previously identified as ssRNA phages that contain coat genes. **Bottom:** contigs previously classified as ssRNA phages that lack coat protein genes.

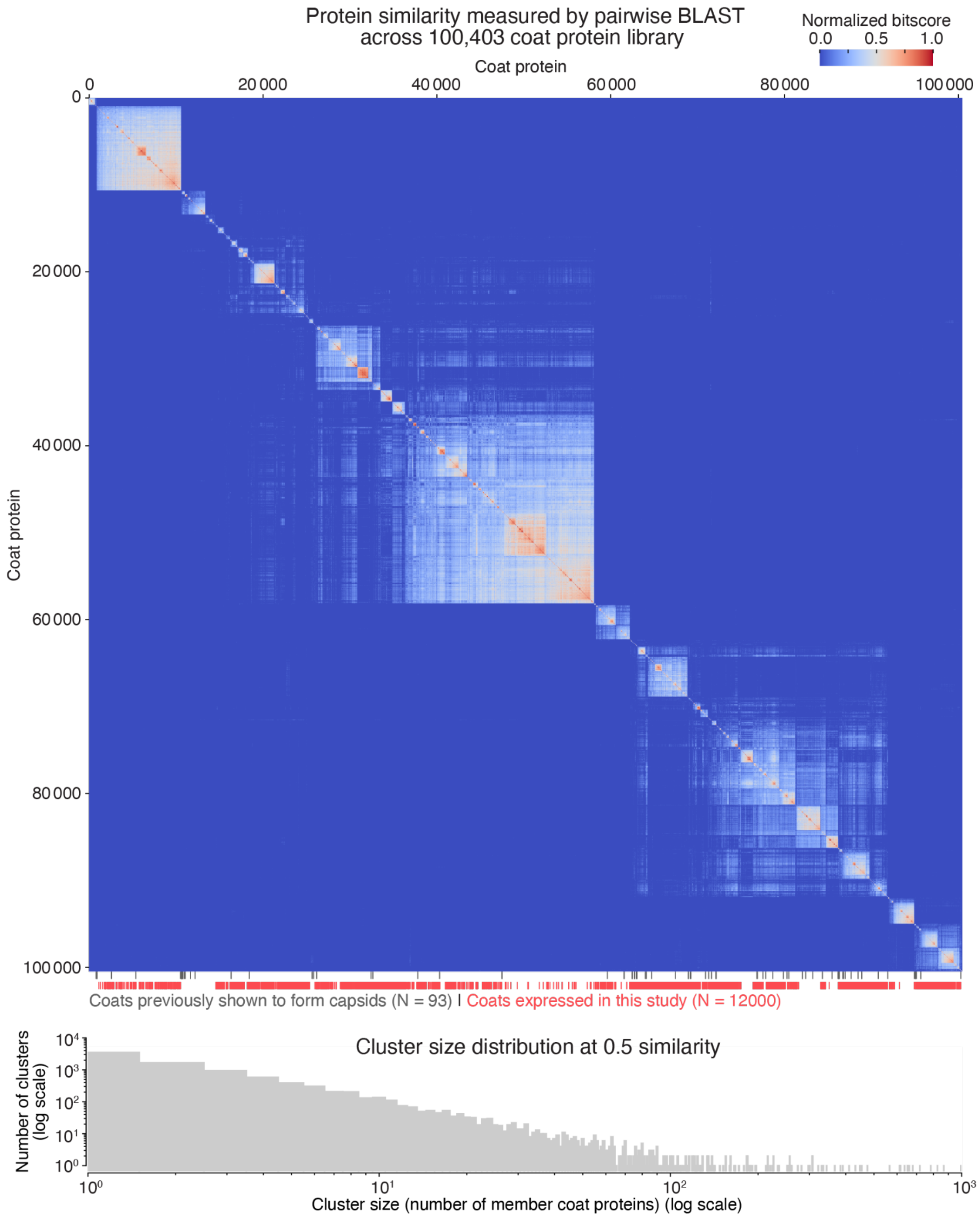


**Fig. S5. Clustered heatmap of all coat protein sequences. Top:** All-vs-all pairwise BLAST bitscores of the 100k-coat library clustered by sequence similarity. Each score was normalized by the maximum self-score of that pair, and the heatmap was clustered using the UPGMA algorithm. We show the location of 12,000 coats that were experimentally expressed in the main text (red ticks) and 93 coats that have been previously shown to form capsids (gray ticks). **Bottom:** Cluster size distribution for the library clustered at 0.5 similarity.

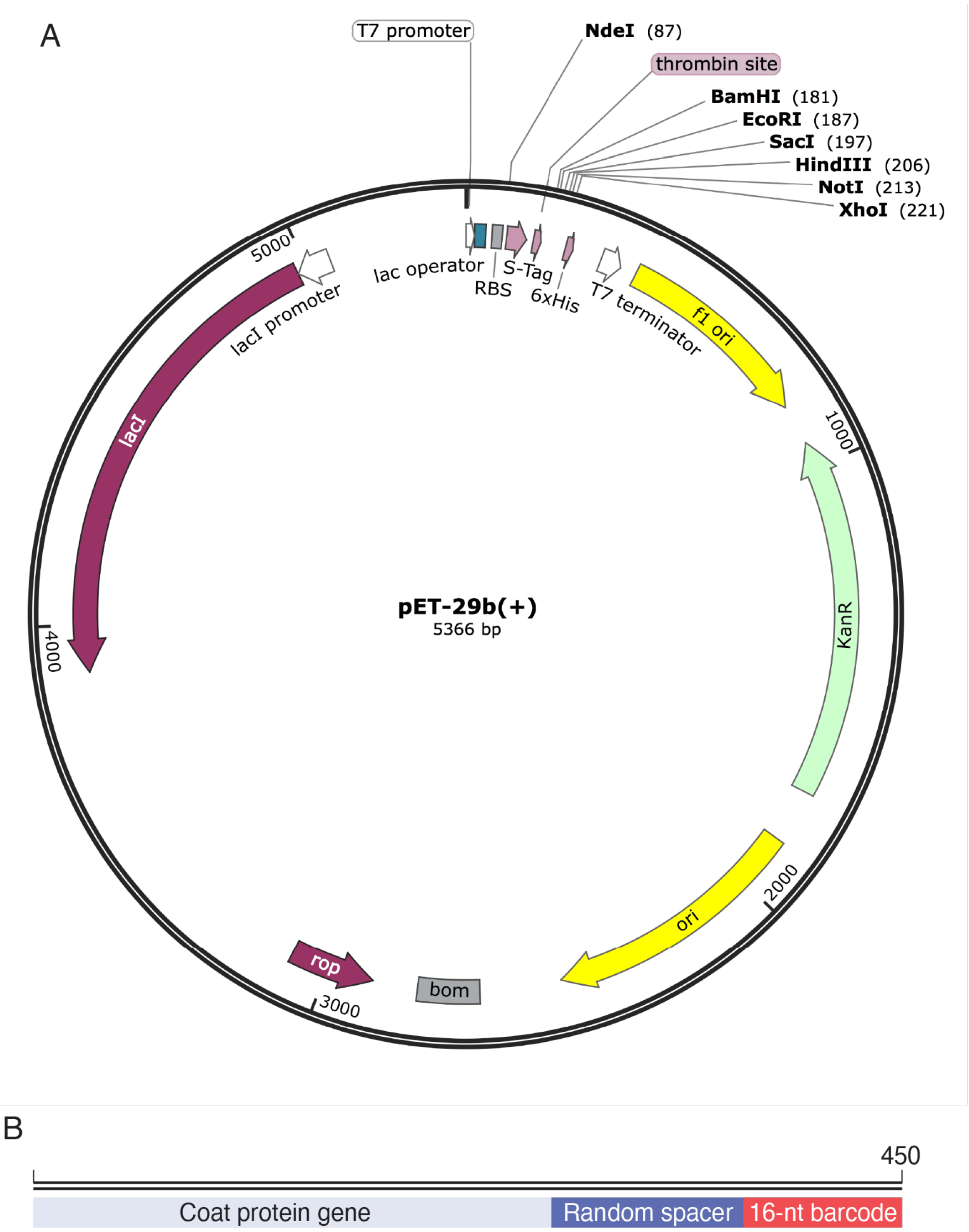


**Fig. S6. Plasmid constructs for the coat-protein library. A.** Map of the pET plasmid vector, provided by Twist Bioscience. The vector contains an IPTG-inducible T7 promoter upstream of the insert sequence. **B.** Map of the insert. Each insert contains a unique coat protein gene (light blue) and a unique 16-nt barcode (red). In addition, the insert contains a variable length random sequence (dark blue) to pad the length of the insert to 450 bp.

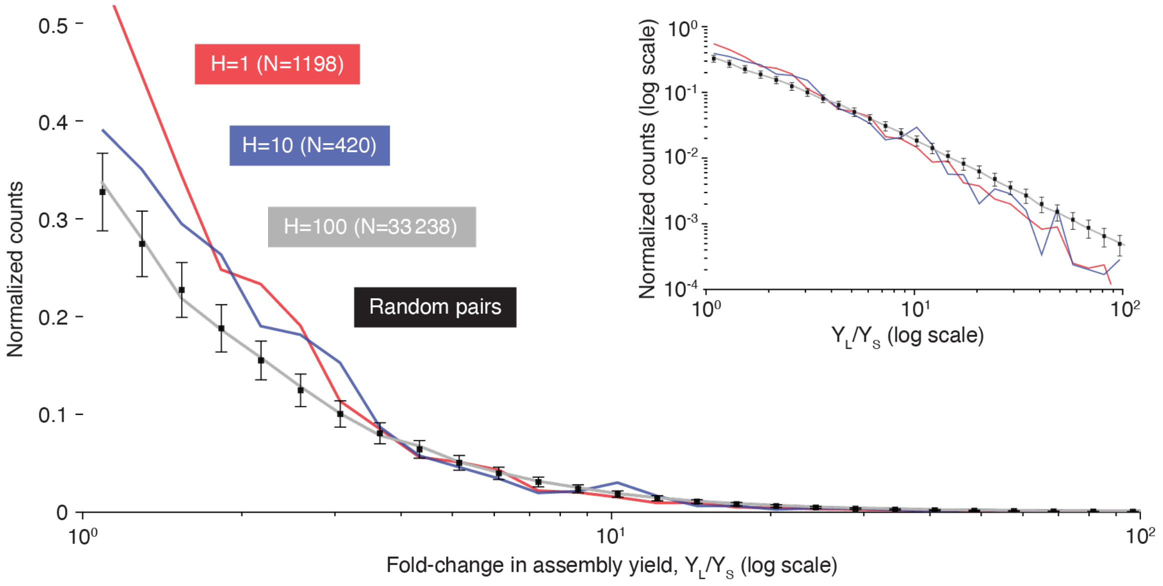


**Fig. S7. The impact of sequence divergence on assembly yield.** In an attempt to understand how the frequency of amino acid substitutions affects capsid assembly yield, we examined all pairs of variants differing by a specific number of substitutions (defined by the Hamming distance, H), and computed the pairwise change in assembly yield (defined as the ratio of the larger yield to the smaller yield in each pair, $Y_L/Y_S$). This analysis captures how much individual (H=1) or multiple (H>1) amino acid substitutions change the assembly yield relative to an otherwise identical counterpart. For three representative values of H (H=1 in red, H=10 in blue, and H=100 in gray), we plotted the density-normalized distribution of $Y_L/Y_S$ ratios observed across all such variant pairs (N denotes the number of pairs). As a control, we also generated 1000 replicates of 1000 randomly selected variant pairs (regardless of sequence similarity), and determined the mean ± standard deviation (shown in black) of their density-normalized $Y_L/Y_S$ distributions. The x-axis is shown on a log-scale to capture the wide range of observed $Y_L/Y_S$ ratios. **Inset:** The same data with the y-axis also shown on a log-scale, to visualize differences at higher $Y_L/Y_S$ ratios. Relative to the control, we observe that the H=1 distribution is significantly enriched for pairs with small changes in assembly yield ($Y_L/Y_S$ ratios less than 3), and depleted in pairs with large changes in yield ($Y_L/Y_S$ ratios greater than 10), with a median $Y_L/Y_S$ ratio of 2.5. The H=10 distribution shows similar trends as the H=1 distribution, with a slightly higher median yield ratio of 3.0. By contrast, the H=100 distribution is indistinguishable from the random distribution, across all yield ratios, with a median of 4.5. While these results suggest that smaller numbers of amino acid substitutions tend to have more modest effects on assembly yield, we note that some H=1 pairs have $Y_L/Y_S$ of 100 or more, indicating that even single substitutions can have dramatic effects on assembly.

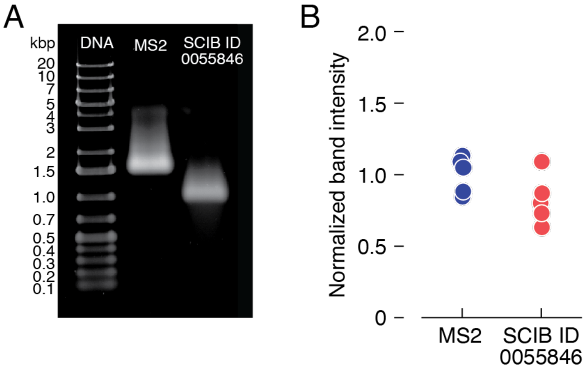


**Fig. S8. Comparing the absolute assembly yield of one isolated variant to that of MS2. A.** We expressed the coat proteins of MS2 and SCIB_ID_0055846 under the same conditions and measured the assembly yield by native agarose gel electrophoresis. We grew cells for 24 h, pelleted the cells, measured the pellet mass, resuspended the pellets in a buffer volume proportional to pellet mass, lysed the cells by sonication, treated the lysate with DNase I and RNase A, performed electrophoresis, and imaged the gel with ethidium bromide to stain RNA protected within assembled capsids. We performed n = 5 technical replicates by loading each sample 5 times across the gel. Only one of the replicates is shown. **B.** We integrated the intensity of the bands in the gel image and normalized these band intensities to the mean band intensity of the MS2 sample.

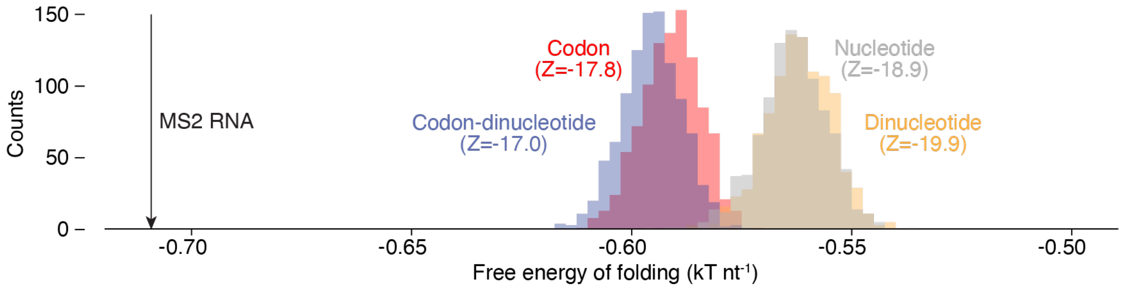


**Fig. S9. Different shuffling strategies give rise to similar MFE Z-scores.** We computed Z-scores for MS2 RNA using different random shuffling procedures, including the standard shuffle that only preserves the frequency of the nucleotides while changing their order (shown in gray), and more complicated shuffling strategies, including those that preserve the dinucleotide frequency (yellow), or the codon frequency within coding regions (red), or both the codon frequency and the dinucleotide frequency (blue). We observed that the different shuffling procedures gave rise to different Z-scores, ranging from -17 to -19.9, but that these differences were relatively small compared to the total magnitude of the Z-scores (only 15% or so). We therefore chose to use the standard shuffling procedure throughout our study.

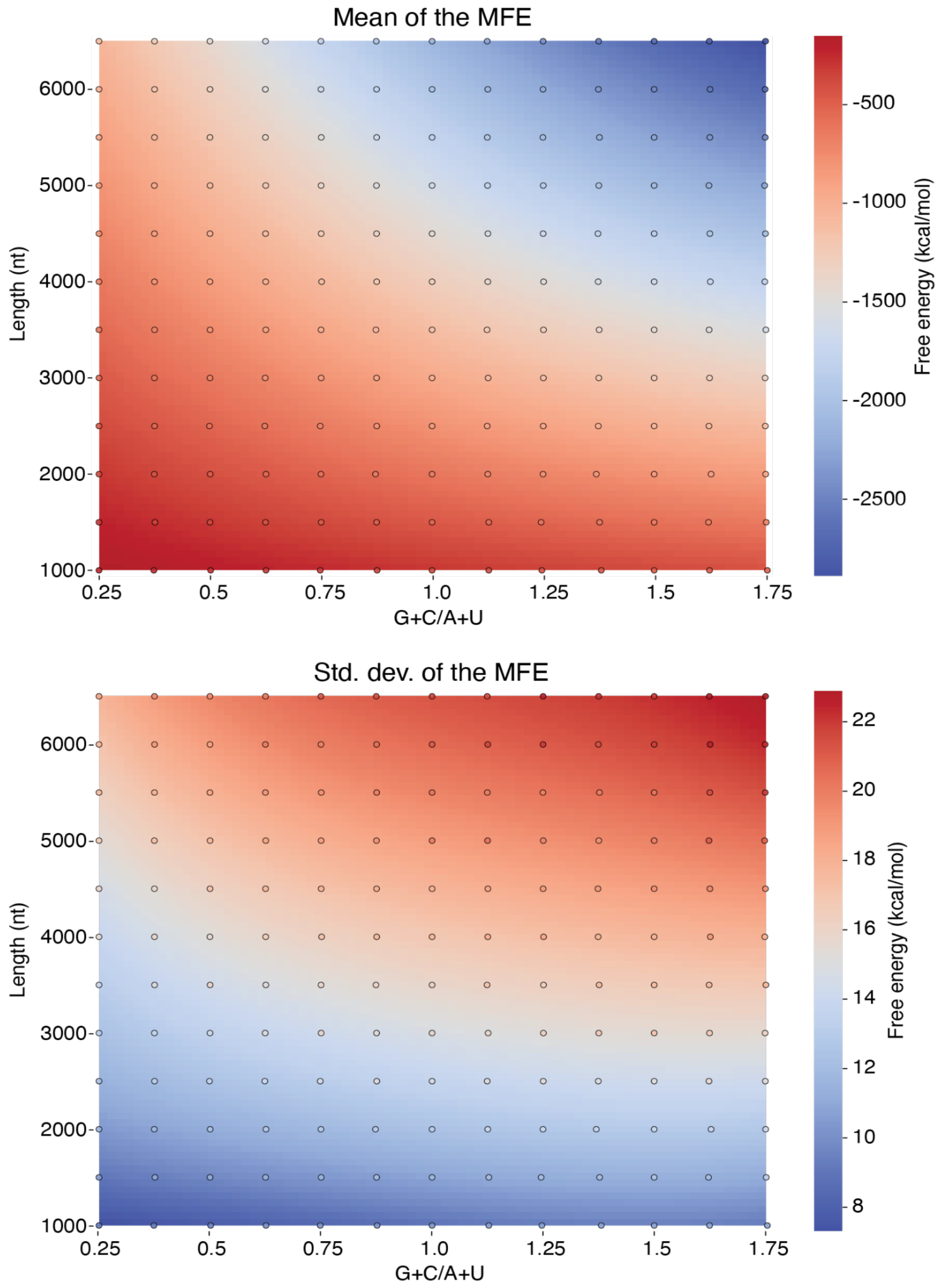


**Fig. S10. Determining Z-scores using a precomputed library of random sequences.** To facilitate Z-score calculations across large datasets, we generated a reference library of shuffled RNA sequences along a grid of defined combinations of sequence length and GC/AU ratio. For each grid point, we calculated MFEs of 1,000 randomly shuffled sequences and computed the mean and standard deviation of the MFE values. We then fit two-dimensional polynomial surfaces of degree four to these values across the parameter space. The resulting surfaces allow interpolation of the expected mean and standard deviation of the MFE for any input sequence within the bounds of the precomputed grid. This approach enables the Z-score of a sequence to be computed directly from that sequence's MFE, length, and GC content, without requiring additional shuffling. **A.** Heatmap showing the estimated mean MFE as a function of length and G+C/A+U ratio. **B.** Heatmap showing the estimated standard deviation of the MFE across the same parameter space. Circles indicate the data points used to fit the surfaces.

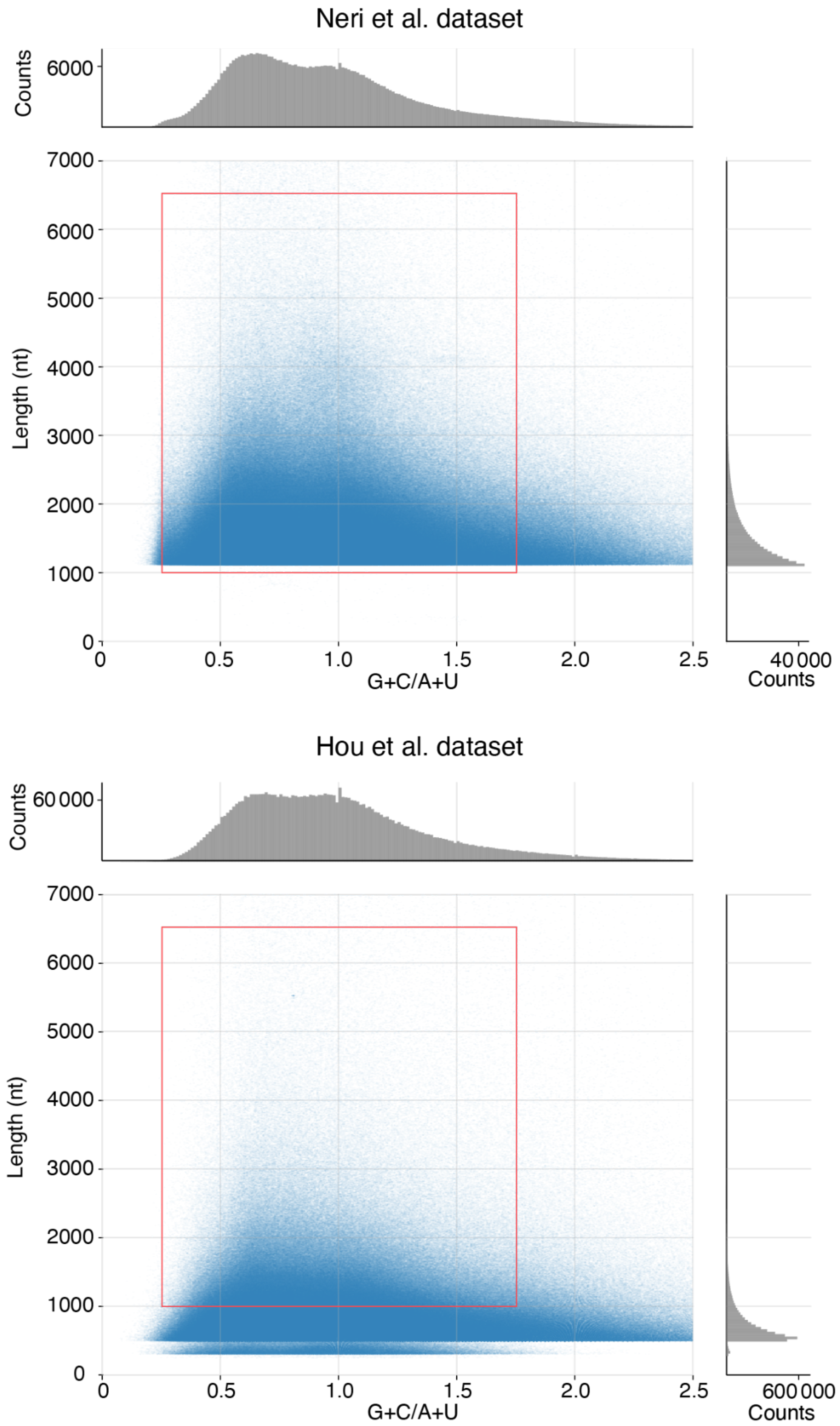


**Fig. S11. Length and G+C/A+U distributions of contigs in the Neri and Hou datasets.** Each contig is plotted as a blue dot. Red boxes show length and G+C/A+U ratios analyzed using RNA folding models. Marginal histograms show the distributions of length (right) and G+C/A+U (top). **Top:** Neri dataset. **Bottom:** Hou dataset.

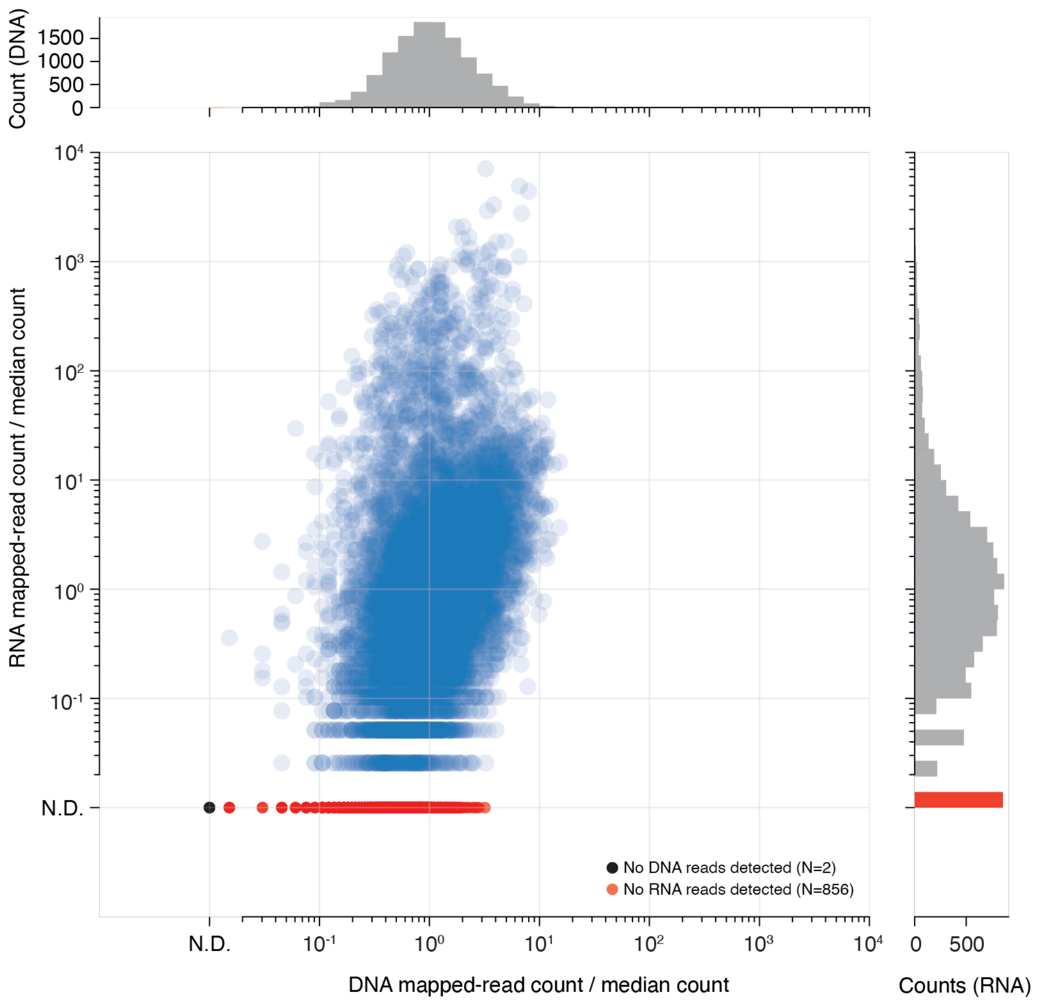


**Fig. S12. Mapped-read counts from DNA and RNA sequencing experiments.** Scatter plot comparing DNA and RNA read counts for each coat-protein barcode, normalized by the median count across all variants in the dataset. The x-axis shows DNA read counts, providing a measure of variant prevalence in the input library. The y-axis shows the RNA read count, a measure of variant prevalence in the packaged RNA. Each point represents a unique coat-protein variant. Variants with no detectable reads in DNA or RNA are plotted as “N.D.” (not detected) and labeled as either black (no DNA counts detected, N=2) or red (no RNA counts detected, N=856). Marginal histograms show the distribution of normalized read counts across the DNA (top) and RNA (right) datasets.

| **Map** | T3 | T4 | D5 |
| --- | --- | --- | --- |
| EMD Identifier | EMD-XXXXX | EMD-XXXXX | EMD-XXXXX |
| PDB Identifier | XXXXX | XXXX | XXXX |
| **Data collection and processing** | | | |
| Magnification | 130,000 | 130,000 | 130,000 |
| Voltage (kV) | 200 | 200 | 200 |
| Electron exposure ($e^-/Å^2$) | 44.4 | 44.4 | 44.4 |
| Defocus range (μm) | -0.5 to - 2.0 | -0.5 to - 2.0 | -0.5 to - 2.0 |
| Pixel size (Å) | 0.886 | 0.886 | 0.886 |
| Symmetry imposed | I1 | I1 | D5 |
| Number of micrographs | 5,279 | 5,279 | 5,279 |
| Number of particles | 287,912 | 48,245 | 10,130 |
| Map resolution (Å, $FSC_{0.143}$) | 2.4 | 2.8 | 3.4 |
| **Refinement** | | | |
| R.m.s. deviations | | | |
| Bond lengths (Å) | 0.010 | 0.004 | 0.003 |
| Bond angles (°) | 1.199 | 0.801 | 0.662 |
| Validation | | | |
| MolProbity score | 1.76 | 2.39 | 1.92 |
| Clashscore | 3.25 | 10.32 | 10.44 |
| Poor rotamers (%) | 5.82 | 7.88 | 0.21 |
| Ramachandran plot | | | |
| Favored (%) | 97.70 | 96.98 | 94.46 |
| Allowed (%) | 2.30 | 3.02 | 5.27 |
| Disallowed (%) | 0.00 | 0.00 | 0.27 |
| CC (volume) | 0.92 | 0.87 | 0.87 |

**Table S1.** Table of cryo-EM data for eliophage capsids. We will provide EMD info upon publication.
